# Rod-shaped tricalbins contribute to PM asymmetry at curved ER-PM contact sites

**DOI:** 10.1101/579128

**Authors:** Patrick C. Hoffmann, Tanmay A. M. Bharat, Michael R. Wozny, Elizabeth A. Miller, Wanda Kukulski

**Affiliations:** Cell Biology Division, MRC Laboratory of Molecular Biology, Francis Crick Avenue, Cambridge CB2 0QH, United Kingdom; Sir William Dunn School of Pathology, University of Oxford, South Parks Road, Oxford OX1 3RE, United Kingdom; Central Oxford Structural and Molecular Imaging Centre, South Parks Road, Oxford OX1 3RE, United Kingdom

**Keywords:** Membrane contact sites, Lipid transfer protein, Endoplasmic reticulum, Plasma membrane, Extended-Synaptotagmin, Tricalbin, Correlative light and electron microscopy, Cryo-focused ion beam milling, Electron cryo-tomography, High-throughput yeast genetics

## Abstract

Lipid flow between cellular organelles occurs via membrane contact sites that form dynamic conduits. Extended-synaptotagmins, known as tricalbins in yeast, mediate lipid transfer between the endoplasmic reticulum (ER) and plasma membrane (PM). How these proteins regulate the membrane architecture to transport lipids across the aqueous space between bilayers remains unknown. Using correlative microscopy, electron cryo-tomography and high-throughput genetics we address this interplay of architecture and function in budding yeast. We find that ER-PM contacts are diverse in protein composition and membrane morphology, not in intermembrane distance. *In situ* cryo-EM of tricalbins reveals their molecular organisation that suggests an unexpected structural framework for lipid transfer. Genetic analysis identifies functional redundancies, both for tricalbin domains and cellular lipid routes, and points to tricalbin function in maintenance of PM asymmetry. These results uncover a modularity of molecular and structural functions of tricalbins, and of their roles within the network of cellular lipid fluxes.

## Introduction

The endoplasmic reticulum (ER) forms a vast membrane network that is structured into thin tubules, densely reticulated regions as well as extended sheet-like cisternae (Nixon-Abell et al., 2016; West et al., 2011). This network interacts with virtually all other organelles through membrane contact sites (MCS) that mediate metabolite exchange and signalling (Valm et al., 2017). At the cell periphery, the ER is in contact with the plasma membrane (PM) by close apposition of 10 to 30 nm (Fernández-Busnadiego et al., 2015; West et al., 2011). Such ER-PM contacts provide spatial control over Ca^2+^ regulation and signalling, and mediate key transfer steps in lipid metabolism (Saheki and De Camilli, 2017). In budding yeast, about one third of the PM is covered by ER (Loewen et al., 2007), which is therefore often referred to as cortical ER (cER). The vast majority of yeast ER-PM contacts are mediated by a set of six proteins from three families, which are conserved between yeast and humans: two VAMP-associated proteins (VAPs), Scs2 and Scs22; TMEM16-like Ist2; and three Extended-Synaptotagmin (E-Syts) orthologues, called tricalbins Tcb1, Tcb2 and Tcb3 (Manford et al., 2012). All six proteins are integral to the ER membrane and bind to the PM by interacting with lipid head groups or other proteins. Proteins of the three families differ in domain organisation as well in the length of the putative linkers bridging the contact site (Gatta and Levine, 2016).

Besides their structural role in membrane bridging, ER-PM proteins contribute additional functionality to the ER-PM interface. The VAPs are interaction hubs for more than 100 proteins, many of which are lipid transfer proteins (LTPs) specific for MCS (Loewen et al., 2003; Murphy and Levine, 2016). The TMEM16 family consists of Cl^-^ channels and lipid scramblases (Brunner et al., 2014; Suzuki et al., 2010). E-Syts are directly capable of transferring lipids between membranes (Saheki et al., 2016; Yu et al., 2016), and absence of tricalbins or E-Syts leads to impaired control of PM homeostasis (Bian et al., 2018; Omnus et al., 2016). Like E-Syts, yeast tricalbins contain a synaptotagmin-like mitochondrial-lipid-binding protein (SMP) domain as well as four or five C2 domains (Giordano et al., 2013; Kopec et al., 2010; Schulz and Creutz, 2004; Toulmay and Prinz, 2012). The lipid-harbouring SMP domain is required for *in vitro* lipid transfer, while at least some of the C2 domains bind to PI(4,5)P_2_ in the PM in a Ca^2+^-dependent manner (Giordano et al., 2013; Schauder et al., 2014; Schulz and Creutz, 2004).

ER-PM contact sites clearly have complex macromolecular architectures with diverse components that contribute to multiple cellular processes. Protein organisation and function are profoundly coupled at these sites, yet detailed understanding on the interplay between protein structure, membrane architecture and contact site function is lacking. Furthermore, the contributions of individual contact site proteins to cell physiology remain difficult to assess, likely due to redundancies (Saheki and De Camilli, 2017; Wong et al., 2017). We have combined correlative light and electron microscopy (CLEM), electron cryo-tomography (cryo-ET) of cryo-focused ion beam (cryo-FIB)-milled cells, and live-cell imaging with high content yeast genetics to unravel the intricate relationship between structure and function of ER-PM contact sites in budding yeast.

## Results

### ER-PM proteins associate with distinct ER membrane shapes, but drive similar intermembrane distances

We first investigated whether the protein families mediating ER-PM contacts are distributed homogenously throughout the cER. We imaged live yeast cells in which we chromosomally tagged pairs of ER-PM proteins with fluorescent proteins (Figure 1). All tagged proteins localized to cER as described before (Loewen and Levine, 2005; Manford et al., 2012; Toulmay and Prinz, 2012; Wolf et al., 2012). We analysed the degree of colocalization among the different pairs by plotting fluorescence intensity profiles along the cell cortex. The paired profiles of Tcb3-mRuby and GFP-Scs2, as well as of Tcb3-mRuby and GFP-Ist2, overlapped to large extents, indicating colocalization within most of the cER (Figure 1A and B). However, in both cases the paired profiles did not completely overlap. Individual peaks of intensity indicated regions at which either of the proteins was enriched relative to the other. In contrast, the intensity profile of Tcb1-GFP overlapped completely with Tcb3-mRuby (Figure 1C). These data suggest partial segregation of the different protein families within the cER, and thus the existence of ER-PM contacts with differential composition.

**Figure 1.**
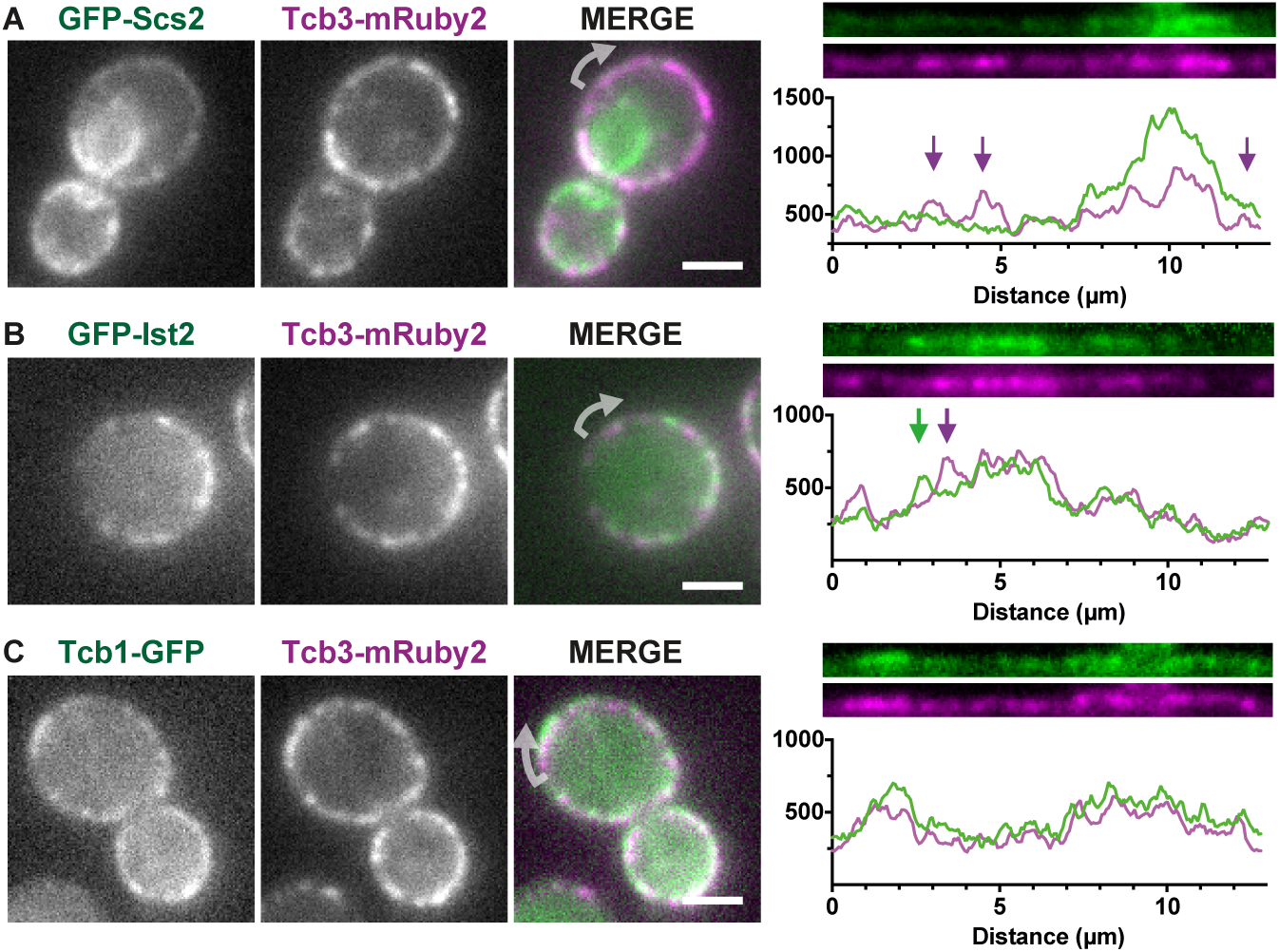
Proteins mediating ER-PM contacts are not distributed homogenously across the cER. Live FM of yeast cells expressing Tcb3-mRuby2 in combination with either (**A**) GFP-Scs2, (**B**) GFP-Ist2 or (**C**) Tcb1-GFP. All proteins are expressed from their endogenous genomic loci. In the merge of the two channels, arrows indicate the starting point of the linearized signals along the mother cell cortex, shown in the right panels, which also show the line profiles of linearized signals (pixel intensity in arbitrary units). Regions in which the signals differ are indicated by arrows. Scale bars: 2 *µ*m.

We next asked whether partial segregation could be linked to differences in ER-PM ultrastructure. We reasoned that local accumulation of individual proteins within the cER might generate functionally specific environments by modulating the ultrastructure. We applied CLEM on resin-embedded yeast cells (Kukulski et al., 2011). In electron tomograms acquired at locations of GFP signals, we visualized the 3D membrane architecture of the ER and the PM correlated with the presence of GFP-Scs2, GFP-Ist2 or Tcb3-GFP (Figure 2A-C). In these tomograms we also found regions of cER that were not correlated with GFP signals, thus indicating absence or very low levels of the GFP-tagged protein (Figure 2A and C, orange arrows). These results corroborate that proteins of the different families partially segregate within the cER. Moreover, some of the cER regions devoid of the protein of interest were immediately continuous with cER regions containing the protein of interest, indicating that segregation also occurs within individual cER subcompartments.

**Figure 2.**
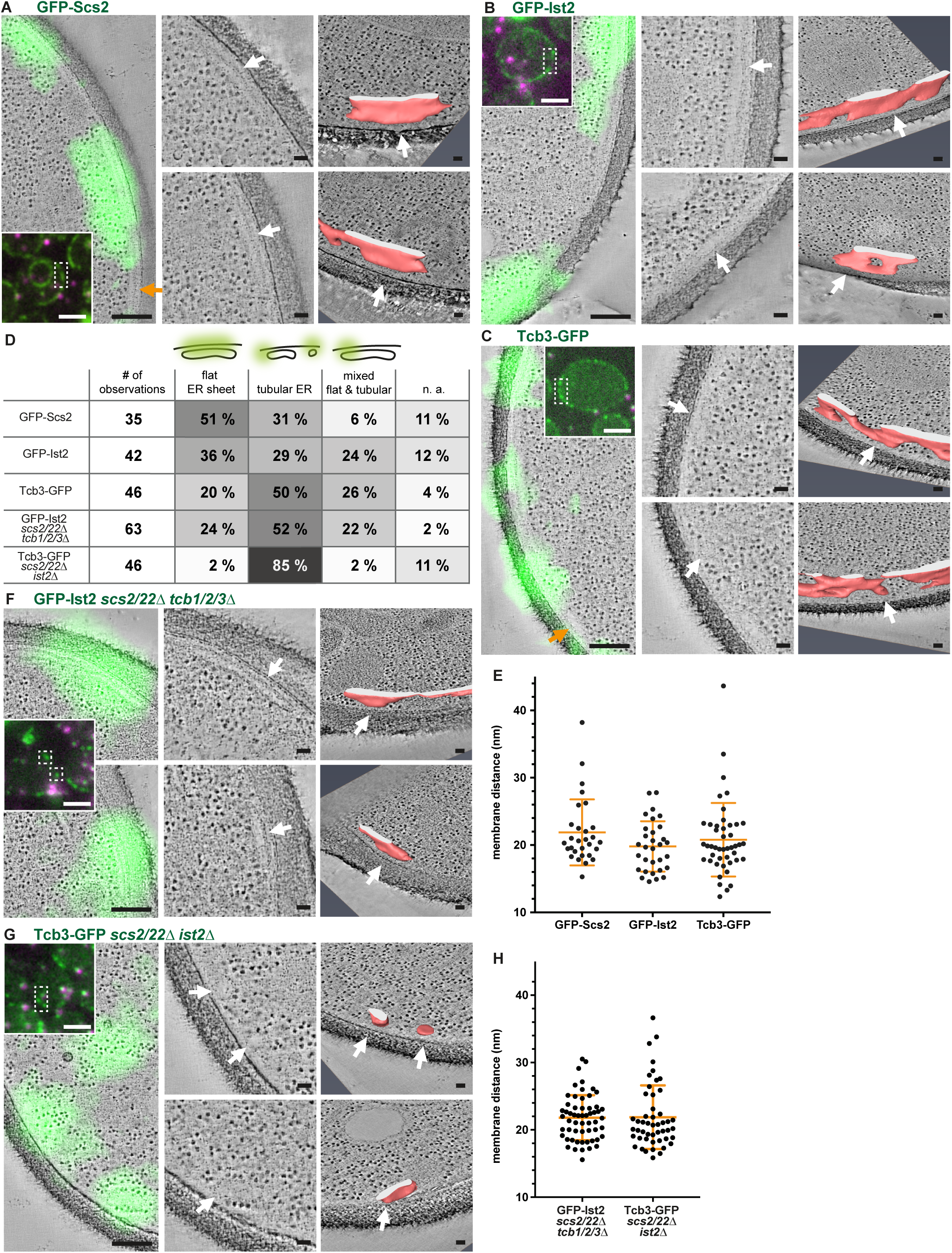
ER-PM proteins associate with distinct ER shapes, but mediate similar intermembrane distances. **A-C, F, G:** CLEM of resin-embedded yeast cells, genotype indicated above panels. **Insets:** FM of resin sections of wild type (A-C) and mutant (F, G) cells expressing GFP-tagged ER-PM proteins (green), fiducial markers are visible in magenta and green. Left images in panels A-C, F and G are virtual slices through electron tomograms, overlaid with the GFP signals transformed according to fiducial-based correlation. Middle panels are magnified views of the same virtual slices, depicting the ER associated with GFP signals. Arrows indicate matching positions with images in right panels, which show virtual slices and segmentation models (rotated for better visibility; red: membrane, white: intersection with boundary of resin section) of the cER associated with GFP signals. **D:** Classification of cER morphologies associated with GFP signals in cells with different genotypes. **E, H:** Distances between cytosolic leaflets of ER and PM, measured at contact sites associated with GFP signals. Orange lines indicate mean and SD. Scale bars: 2 *µ*m in FM images, 200 nm in FM-ET overlays, 50 nm in ET and segmentation panels.

We classified the observed cER shapes into sheet-like flat ER cisternae, tubular ER of high membrane curvature, and ER consisting of both flattened and tubular segments (mixed curvature) (Figure 2D). GFP-Scs2 and GFP-Ist2 predominately associated with flat cER sheets (51% and 36% of observations, respectively). Tcb3-GFP localized predominantly to tubular ER (50% of observations). At sites associated with presence of the different GFP-tagged proteins, we measured the distances between the cytosolic leaflets of the PM-facing ER membrane and the PM (see Materials and Methods), and found that they were similar (Figure 2E). The average distance for GFP-Scs2 containing contact sites was 21.9 nm (SD 4.9 nm, N = 29), for GFP-Ist2 19.8 nm (SD 3.7 nm, N = 31) and for Tcb3-GFP 20.8 nm (SD 5.5 nm, N = 45).

Given the high degree of colocalization between the different bridging proteins (Figure 1), the observed cER architectures likely contained not only the GFP-tagged protein, but also some or all of the other bridging proteins. To assess the impact of the different proteins separately, we applied CLEM to cells expressing only one type of bridging protein. In the absence of VAPs and tricalbins (*scs2/22*Δ *tcb1/2/3*Δ cells), GFP-Ist2 was associated more often with tubular ER (52% of observations) than with sheets (24% of observations) (Figure 2D, F). Tcb3-EFGP-containing cER in absence of VAPs and Ist2 (*scs2/22*Δ *ist2*Δ cells) was predominantly tubular (85% of observations) (Figure 2D, G). The intermembrane distances were similar for Tcb3-GFP in *scs2/22*Δ *ist2*Δ cells (average 21.9 nm, SD 4.7 nm, N = 49,) and for GFP-Ist2 in *scs2/22*Δ *tcb1/2/3*Δ cells (average 21.8 nm, SD 3.3 nm, N = 57)(Figure 2H).

These results show that the localization of different bridging proteins correlates with differences in cER shape. In particular, Tcb3 displays a distinct preference for contacts between PM and tubular cER of high membrane curvature. In contrast, the intermembrane distances mediated by different bridging proteins are very similar. Taken together, these data indicate that differences in bridging protein composition of ER-PM contacts translate into differences in cER membrane curvature, rather than differences in inter-organelle distance.

### Curvature of tricalbin-mediated cortical ER is dictated by the tricalbin transmembrane domain

We next investigated the preference of tricalbins for high membrane curvature. ER tubulation is facilitated by reticulons, and absence of these proteins results in almost complete lack of tubular ER (Voeltz et al., 2006; West et al., 2011). Reticulons mediate membrane curvature through a hairpin structure inserted into the ER membrane (Hu et al., 2008; 2009; Voeltz et al., 2006). The ER-integral domain of E-Syts is suggested to form a similar hairpin (Giordano et al., 2013). We first tested how localization of Tcb3 in live yeast cells depends on reticulons. We imaged Tcb3-GFP in relation to Sec63-RFP, a widely used ER marker, expressed from a plasmid (Metzger et al., 2008). In wild type cells, Tcb3-GFP and Sec63-RFP colocalized at the cER, albeit with some distinct areas of reduced correspondence (Figure 3A). In the absence of the two major reticulon-like proteins Rtn1 and Yop1 (*rtn1*Δ *yop1*Δ cells), Tcb3-GFP and Sec63-RFP showed almost no colocalization. Although both proteins still localized to the cell cortex, Sec63-RFP localized as extended sheet-like signals, as observed before (Voeltz et al., 2006), whereas Tcb3-GFP localized exclusively as punctae (Figure 3B). We observed the same localization pattern for Tcb2-GFP and plasmid-encoded Tcb3-GFP in combination with Sec63-RFP (Supplementary Figure S1). In contrast, GFP-Ist2 and GFP-Scs2 entirely overlapped with Sec63-RFP in *rtn1*Δ *yop1*Δ cells (Supplementary Figure S1). When we performed CLEM on Tcb3-GFP *rtn1*Δ *yop1*Δ cells, we found that Tcb3-GFP localized almost exclusively to highly curved tubules at the cell cortex (Figure 3C and E). The same cells also contained extended, smooth cER sheets that were not associated with Tcb3-GFP (Figure 3D). These data indicate that the existence of highly curved cER tubules, which accommodate Tcb3-GFP, does not require reticulons.

**Figure 3.**
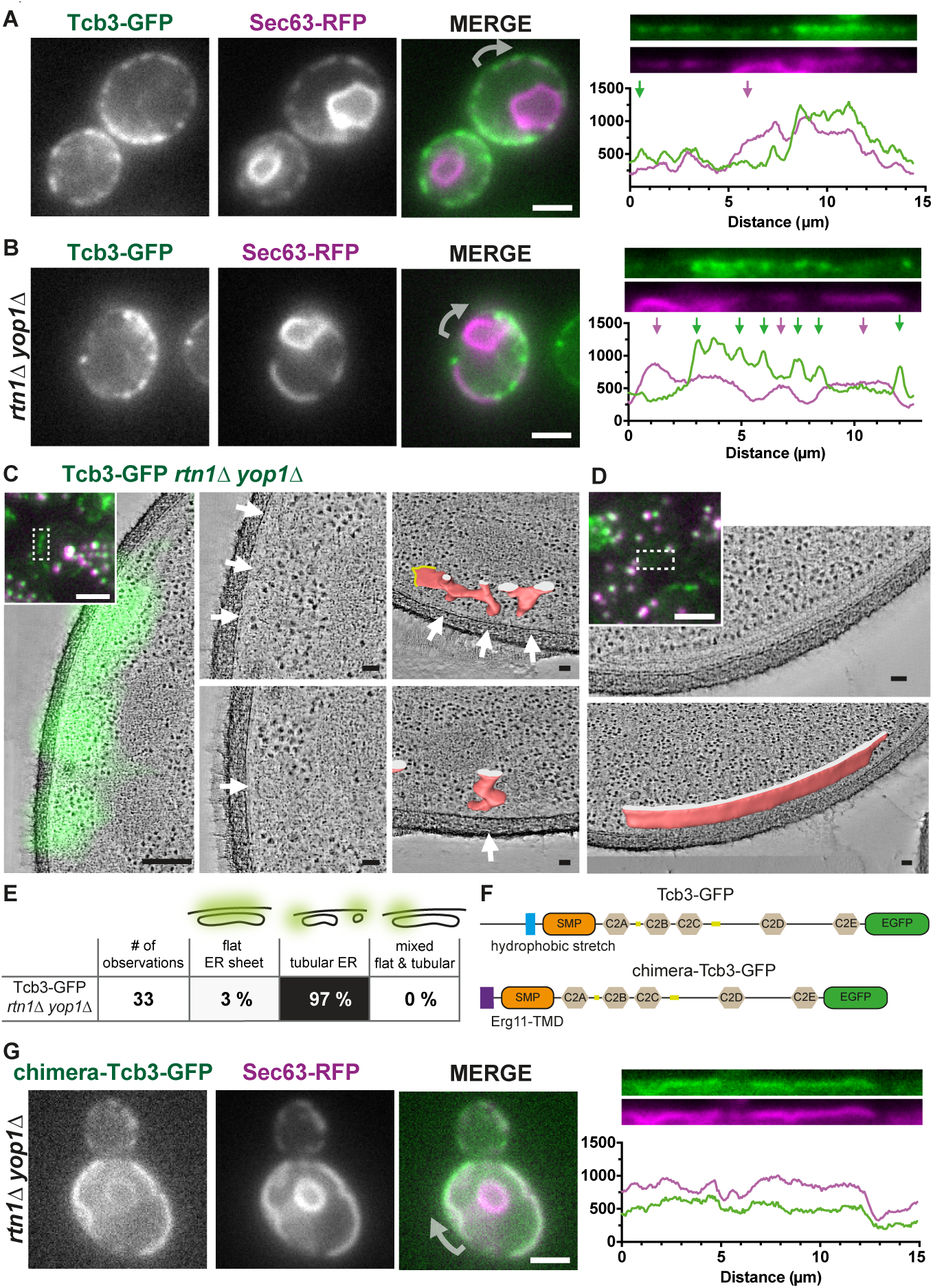
The ER membrane curvature at Tcb3-mediated contact sites is dictated by the Tcb3 transmembrane domain. **A, B:** Live FM of wild type (A) and *rtn1*Δ *yop1*Δ (B) yeast cells expressing Tcb3-GFP and Sec63-RFP. In the merge of the two channels, arrows indicate the starting point of the linearized signals along the mother cell cortex, shown in the right panels, which also show the line profiles of linearized signals (pixel intensity in arbitrary units). Arrows indicate regions in which the two signals segregate. **C, D:** CLEM of resin-embedded *rtn1*Δ *yop1*Δ cells expressing Tcb3-GFP. **Insets**: FM of resin sections (GFP in green, fiducial markers in green and magenta). White dashed box corresponds to the area shown in the underlying virtual slice through electron tomogram, in B overlaid with the Tcb3-GFP signal transformed according to fiducial-based correlation. **C:** Middle images are magnified views of the virtual slice shown in left panel, depicting cER associated with Tcb3-GFP. Arrows indicate matching positions with images in right panel, which show virtual slices and segmentation models (rotated for better visibility; red: membrane, white: intersection with boundary of resin section, yellow: segmentation not continued) of the cER associated with Tcb3-GFP. **D:** Extended cortical sheets devoid of Tcb3-GFP signals, lower panel shows segmentation model (rotated for better visibility; red: membrane, white: intersection with boundary of resin section). **E:** Classification of the ER membrane morphology associated with Tcb3-GFP in *rtn1*Δ *yop1*Δ cells. **F:** Tcb3 domain structure. In chimera-Tcb3-GFP, the N-terminal part (1-247aa) including the transmembrane domain is replaced with the ER signal sequence and transmembrane domain (1-55aa) of ER-resident protein Erg11. **G:** Live FM of *rtn1*Δ *yop1*Δ cells expressing plasmid-encoded Sec63-RFP and chimera-Tcb3-EFGP. In the merge of the two channels, arrow indicates the starting point of the linearized signals along the mother cell cortex, shown in the right panel, which also shows the line profile of linearized signals (pixel intensity in arbitrary units). Scale bars: 2 *µ*m in FM, 200 nm in FM-ET overlay, 50 nm in ET and segmentations.

To test the role of the Tcb3 transmembrane domain in the segregation of Tcb3-GFP to cER tubules, we replaced the sequence predicted to encode the hairpin with a single transmembrane alpha-helix from the ER-resident protein Erg11 (Monk et al., 2014). We refer to this construct as chimera-Tcb3-GFP (Figure 3F). In *rtn1*Δ *yop1*Δ cells expressing plasmid-encoded Sec63-RFP, chimera-Tcb3-GFP colocalized extensively with Sec63-RFP in a sheet-like pattern, distinctive from the punctate localization of wild type Tcb3-GFP (Figure 3G). These results indicate that the transmembrane domain of Tcb3 is responsible for localization of Tcb3-GFP to highly curved cER membranes.

In summary, these results show that tricalbins have a clear preference for curved membranes. In wild-type cells, reticulons are known to generate significant curvature of the cER, likely facilitating tricalbin accumulation there. In the absence of reticulons, tricalbins become restricted to small subdomains of residual curvature, devoid of the translocon component Sec63. The N-terminal transmembrane domain of Tcb3 is responsible for this segregation, and possibly even drives curvature directly in these regions.

### Tricalbins functionally interact with lipid sensing, transfer and biosynthesis pathways

Having defined a curvature localization domain of Tcb3, we next wanted to test the functional importance of this domain. Deletion of all three tricalbins causes no significant phenotype (Creutz et al., 2004; Manford et al., 2012; Toulmay and Prinz, 2012), possibly due to redundancies. Therefore, to reveal useful phenotypes that might be complemented by the curvature-insensitive form of Tcb3 (Figure 4A), we performed a synthetic genetic array (SGA) using a *tcb1/2/3*Δ query strain crossed to the haploid gene deletion and DAmP (decreased abundance by mRNA perturbation) collections (*genex*Δ) (Schuldiner et al., 2005; Tong and Boone, 2007) (Figure 4B). We scored growth phenotypes by measuring colony size of quadruple (*tcb1*Δ *tcb2*Δ *tcb3*Δ *genex*Δ) deletion mutants and calculating the log2 ratio of query colony size relative to control strains (see Materials and Methods). We considered a log2 ratio smaller than −1, corresponding to a 50% reduction in colony size, to be a hit. 724 genes showed negative synthetic interactions with *tcb1/2/3*Δ (Figure 4C). We validated selected hits using serial dilution growth assays (Figure 4D). A large number of genes showing synthetic negative interactions corresponded to components involved in cellular lipid metabolism and membrane biology. Among these hits were genes encoding proteins previously characterised to function at MCS, including the nuclear-vacuolar junction proteins Nvj1 and Vac8, and components of the ER-mitochondrial-encounter structure (ERMES) Mdm34 and Mdm10 (Figure 4C, top panel) (Kornmann et al., 2009; Pan et al., 2000). These hits suggest that Tcb3 function is partially redundant with functions occurring at other MCS, consistent with emerging models of lipid flow by parallel routes. We also identified genes involved in autophagy, more specifically autophagophore formation (Figure 4C, second panel). One of these genes, *atg2*, has been suggested to encode a lipid transfer protein that shares homology with Vps13 (Gao and Yang, 2018). These hits suggest a role for lipid transfer via tricalbins in generating autophagosomal membranes. Finally, we found all core components of the RIM101 pathway, which is involved in sensing alkaline pH and disturbance of PM lipid asymmetry (Ikeda et al., 2008). These hits suggest a functional link between tricalbins and lipid distribution across the PM (Figure 4C, third panel).

**Figure 4.**
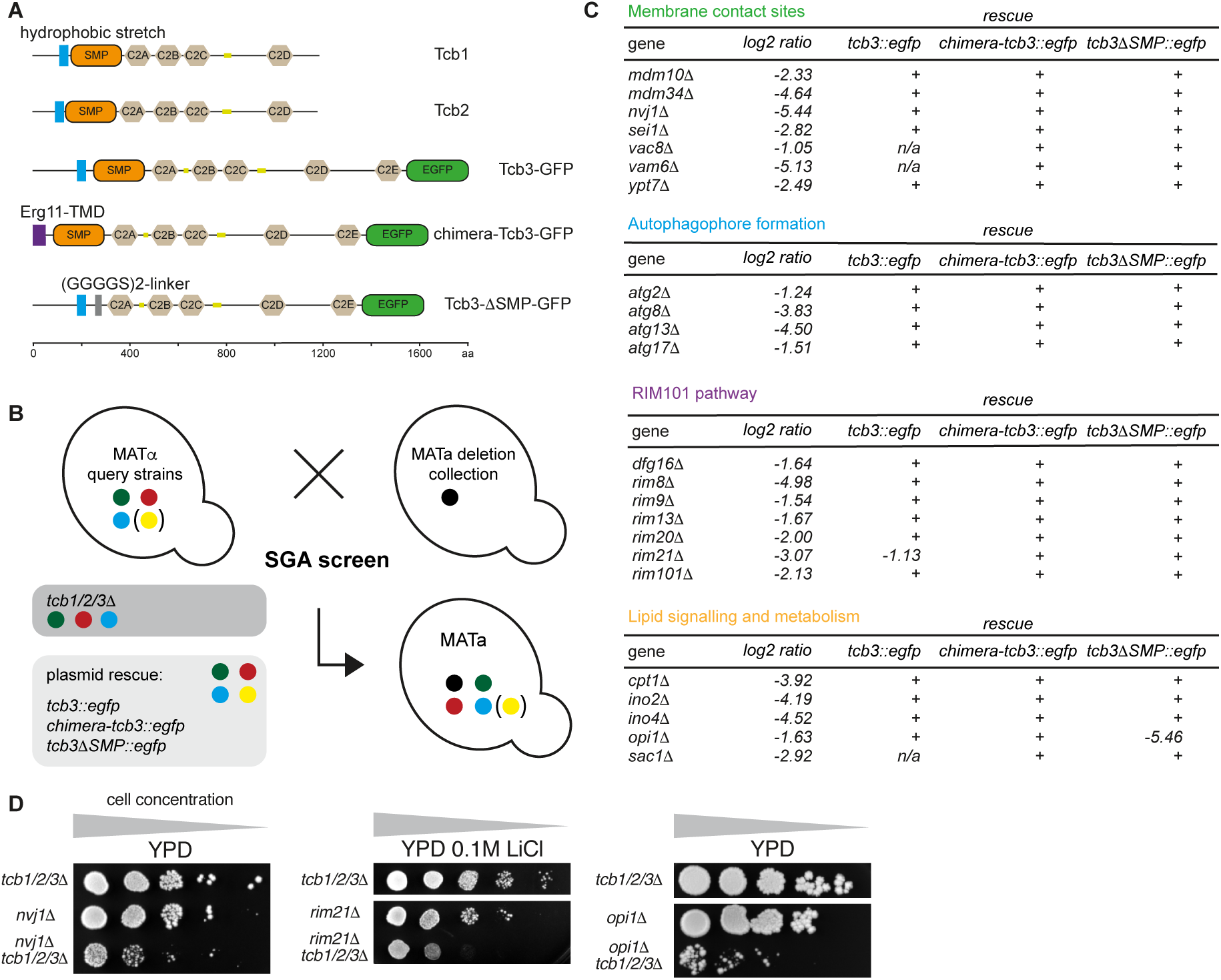
Synthetic genetic interactions of tricalbins. **A:** Sequence and domain structure of tricalbins and rescue constructs; which are plasmid–encoded wild type Tcb3-GFP and mutants with ER anchoring domain replaced (chimera-Tcb3-GFP), or SMP domain replaced (Tcb3-ΔSMP-GFP). **B:** Schematic overview of SGA screen: The query strains, *tcb1/2/3*Δ or *tcb1/2/3*Δ with rescue plasmid, were crossed to the yeast gene deletion and DAmP collections. Haploid mutants containing all deletion markers were selected on appropriate selection media. Colony size was scored and compared to mutants with random distribution of gene deletions. **C:** Negative genetic interactions grouped according to function: membrane contact sites, autophagophore formation, Rim101 pathway, and lipid signalling and metabolism. **D:** Validation of selected synthetic interactions by serial dilution growth assays on various substrates, comparing growth of query strain *tcb1/2/3Δ* to single and quadruple deletions.

We next asked which genetic interactions could be rescued by expression of plasmid-encoded chimera-Tcb3-GFP, the curvature-insensitive form of Tcb3 (Figure 4A). We found that all interactions with genes of the RIM101 pathway were rescued by chimera-Tcb3-GFP to a similar extent as when wild type Tcb3-GFP was expressed from the same plasmid. Moreover, most interactions from the cohorts of MCS and autophagy genes were also fully rescued by chimera-Tcb3-GFP (Figure 4C). In summary, most Tcb3-specific phenotypes were rescued by Tcb3 lacking a curvature-sensitive domain.

The SMP domain of E-Syts is necessary for lipid transfer and is thus considered a major functional domain of E-Syts and Tricalbins (Saheki et al., 2016; Yu et al., 2016). We therefore tested its importance for Tcb3 function by rescue of synthetic interactions in *tcb1/2/3*Δ cells. We replaced the SMP domain in Tcb3-GFP by a linker sequence and expressed it from a plasmid (Figure 4A). Like chimera-Tcb3-GFP, the ΔSMP construct rescued the majority of hits. One exception was *opi1*Δ, whose interaction with *tcb1/2/3*Δ was rescued by wild type Tcb3-GFP and chimera-Tcb3-GFP, but not by Tcb3-ΔSMP-GFP (Figure 4C, bottom panel). Opi1 is a transcriptional repressor of a large number of lipid biosynthetic genes (Wagner et al., 1999). In summary, we have uncovered genetic interactions that indicate potential involvement of tricalbins in specific cellular lipid distribution routes, in particular PM bilayer asymmetry. While these functions depend on the presence of at least one tricalbin in addition to VAPs and Ist2, they neither require this tricalbin to have its curvature-sensitivity nor its SMP domain that presumably mediates lipid transfer.

### The protein organization of tricalbin-mediated contact sites responds to Ca^2+^

The structural organisation of LTPs between two membranes is key to their function (Reinisch and De Camilli, 2016). We therefore set out to visualize tricalbins *in situ* using recent advances in cellular cryo-ET that allow imaging of macromolecular structures within cells (Beck and Baumeister, 2016). We vitrified cells in which ER-PM contacts were formed exclusively by the tricalbins (*scs2/22*Δ *ist2*Δ cells), and subjected these cells to thinning by cryo-FIB milling and to cryo-ET (Marko et al., 2007). In resulting tomograms of cortical cell regions, we found tubular cisternae of cER (Supplementary Figure S2A, B), in agreement with our findings from CLEM on resin-embedded cells (Figure 2G). In addition, the cryo-ET data revealed densities forming bridges between ER and PM (Figure 5A, white arrows). These densities were distributed at irregular intervals and were often positioned between the membranes in a non-perpendicular orientation. Many of the densities appeared rod-like. In some cases, the ER membrane was locally buckled towards the PM at the location of the bridging density (Figure 5A, top white arrow). Since these cells did not express VAPs and Ist2, we conclude that the majority of the bridging densities correspond to tricalbins. Occasionally, patches of PM were coated with a layer of density, reminiscent of a protein coat (Figure 5A, orange arrow).

**Figure 5.**
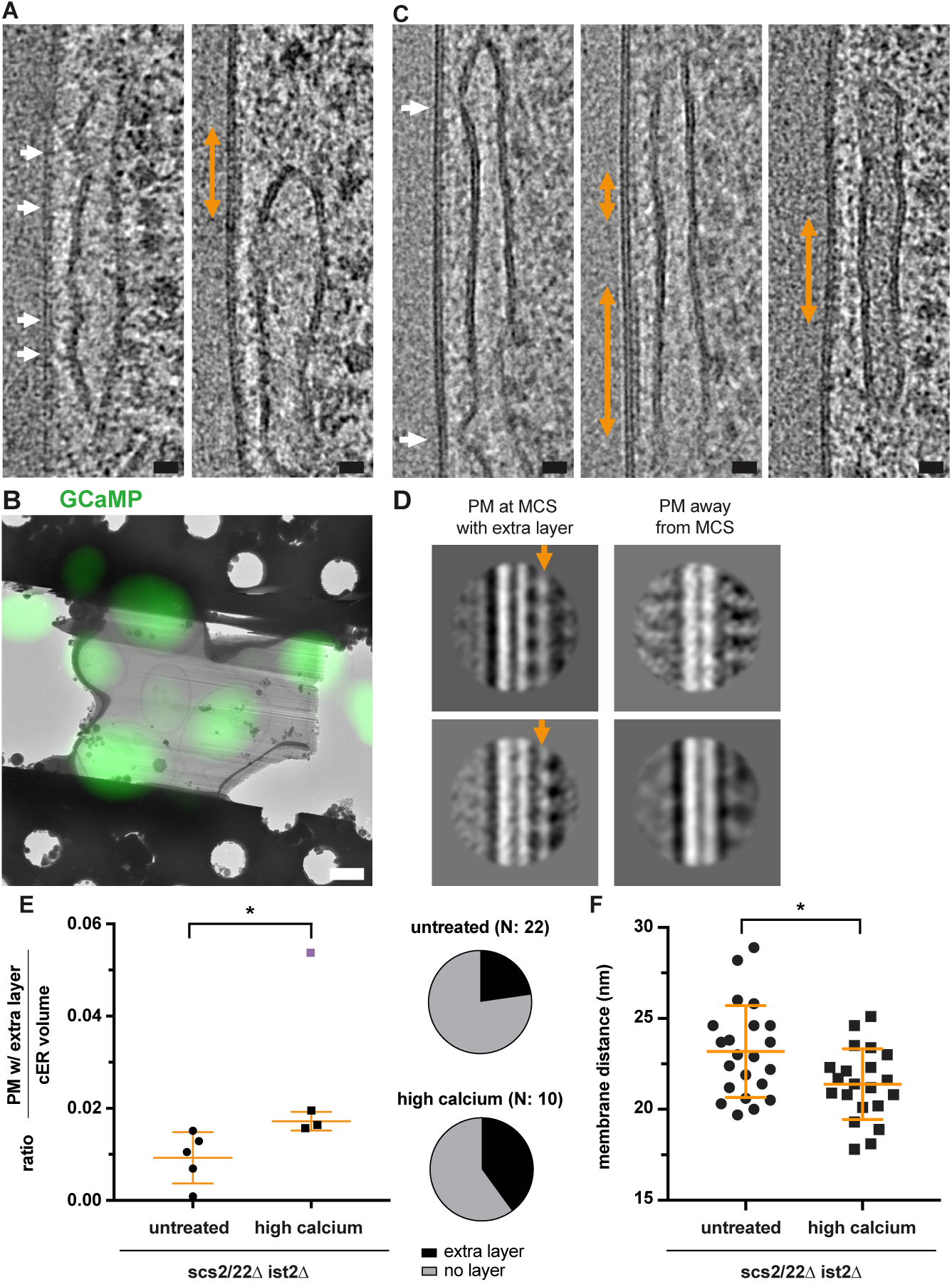
Protein organization of tricalbin-mediated membrane contact sites depends on Ca^2+^. **A:** Virtual slices through electron cryo-tomogram of cryo-FIB milled *scs2/22*Δ *ist2*Δ cells. White arrows indicate densities bridging ER and PM. Orange arrow indicates density layer on PM. **B:** Overlay of cryo-EM image of FIB-milled lamella and cryo-FM image showing GCaMP signal of cells prior to FIB-milling. Cells were treated with 200 mM CaCl_2_ prior to vitrification. **C:** Virtual slices through electron cryo-tomograms of cryo-FIB milled *scs2/22Δ ist2Δ* cells displaying strong GCaMP signals. White arrows indicate buckling of the ER towards the PM. Orange arrows indicate density layer on PM. The left and middle images are from the same tomogram. **D:** Subtomogram averages of PM area at ER-PM contact with density layer (indicated by orange arrow), compared to PM area not in contact with cER. The extracellular PM leaflet is facing left, the cytosolic leaflet right. Top and bottom averages are from layer shown in C middle and right panel, respectively. **E:** Ratio of PM coated with extra density layer / cER volume, for untreated *scs2/22Δ ist2*Δ cells and *scs2/22*Δ *ist2Δ* cells with high GCaMP signals. The highest value data point (purple) in the latter data was treated as outlier and excluded from significance test (P=0.0312). Pie charts indicate the fraction of electron cryo-tomograms in which extra density layers were observed. **F:** Distances between ER and PM in untreated *scs2/22*Δ *ist2*Δ cells and in *scs2/22*Δ *ist2*Δ cells with high GCaMP signals (P=0.0126). Orange lines indicate mean and SD in E and F. Scale bars: 20 nm in A and C, 2 *µ*m in B.

E-Syts are regulated by Ca^2+^, which impacts both their lipid transfer function and their capacity to bind to the PM (Bian et al., 2018; Giordano et al., 2013). We reasoned that elevated cytosolic Ca^2+^ levels might trigger functionally important conformational changes that would be directly visible in *scs2/22*Δ *ist2*Δ cells by cryo-ET. To monitor cytosolic Ca^2+^ concentration, we expressed a GFP-Calmodulin-fusion protein (GCaMP) as a fluorescent indicator (Carbó et al., 2017). We treated cells with 200 mM CaCl_2_ and imaged them by live FM. Shortly after extracellular Ca^2+^ was raised, a subpopulation of cells displayed a strong fluorescent signal, which returned to background levels within seconds, indicating transiently elevated cytosolic Ca^2+^ (Supplementary Figure S2C) (Carbó et al., 2017). To target cells with high levels of cytosolic Ca^2+^ for cryo-ET, we used cryo-FM prior to cryo-FIB milling (Ader et al., 2019). This strategy allowed us to identify cells with strong GCamP signals, after they had been treated with 200 mM CaCl_2_, deposited on EM grids and vitrified (Figure 5B). In the resulting tomograms, we found ER-PM contacts similar to those in untreated cells, including cER membrane patches locally buckled towards the PM (Figure 5C, white arrows). We noticed that the PM of several ER-PM contacts was coated with a layer similar in appea rance to what we observed in untreated cells (Figure 5C, orange arrows). The occurrence of this layer was more frequent in cells with high Ca^2+^ (4/10 tomograms) than in untreated cells (5/22 tomograms). Furthermore, the layer extended over a larger fraction of the cER in cells with high Ca^2+^ (Figure 5E). To better resolve details of this layer, we performed subtomogram averaging, previously used to investigate protein coating of the PM (Bharat et al., 2018) (see Materials and Methods). The resulting subtomogram averages showed an on average 5.2 nm (SD 0.5 nm, N = 4) thick density layer on the cytoplasmic leaflet of the PM (Figure 5D, orange arrows and Supplementary Figure S2H), which was not present on PM areas outside of the contact site. Furthermore, cells with high Ca^2+^ displayed shorter intermembrane distances than untreated cells (average distances: 21.4 nm, SD 1.9 nm, N = 21 for high Ca^2+^, and 23.2 nm, SD 2.5 nm, N = 22 for untreated)(Figure 5F). At the locations where the ER membrane was buckled towards the PM, the two membranes appeared locally closer than within the rest of the contact site (Supplementary Figure S3A). The membrane distances at these locations were also shorter in cells with high Ca^2+^ than in untreated cells (average distances: 14.2 nm SD 1.4 nm, N=19 for high Ca^2+^; 15.2 nm, SD 1.0 nm, N = 19 for untreated, Supplementary Figure S3B).

In summary, our cryo-ET analysis indicates that tricalbin-mediated contact sites contain rod-like densities that bridge the two membranes. In addition, within the contact site areas, the PM partially displays a density layer reminiscent of a protein coat. This layer is more frequent and extended when cytosolic Ca^2+^ is high.

### *In situ* structural analysis of Tricalbin-3

We next investigated the structure of the densities bridging the two membranes. To analyse a large number of densities likely corresponding to a single tricalbin isoform, we deleted 5 of the 6 bridging proteins (5Δ cells; *scs2/22*Δ *ist2*Δ *tcb1/2*Δ), and controlled Tcb3-GFP expression by a galactose-inducible promoter. By live FM, we confirmed that induction by galactose was required to localize Sec63-RFP as well as Tcb3-GFP to the cell cortex in the 5Δ cells (Figure 6A). This experiment verified that overexpressed Tcb3-GFP is sufficient for formation of ER-PM contacts in 5Δ cells.

**Figure 6.**
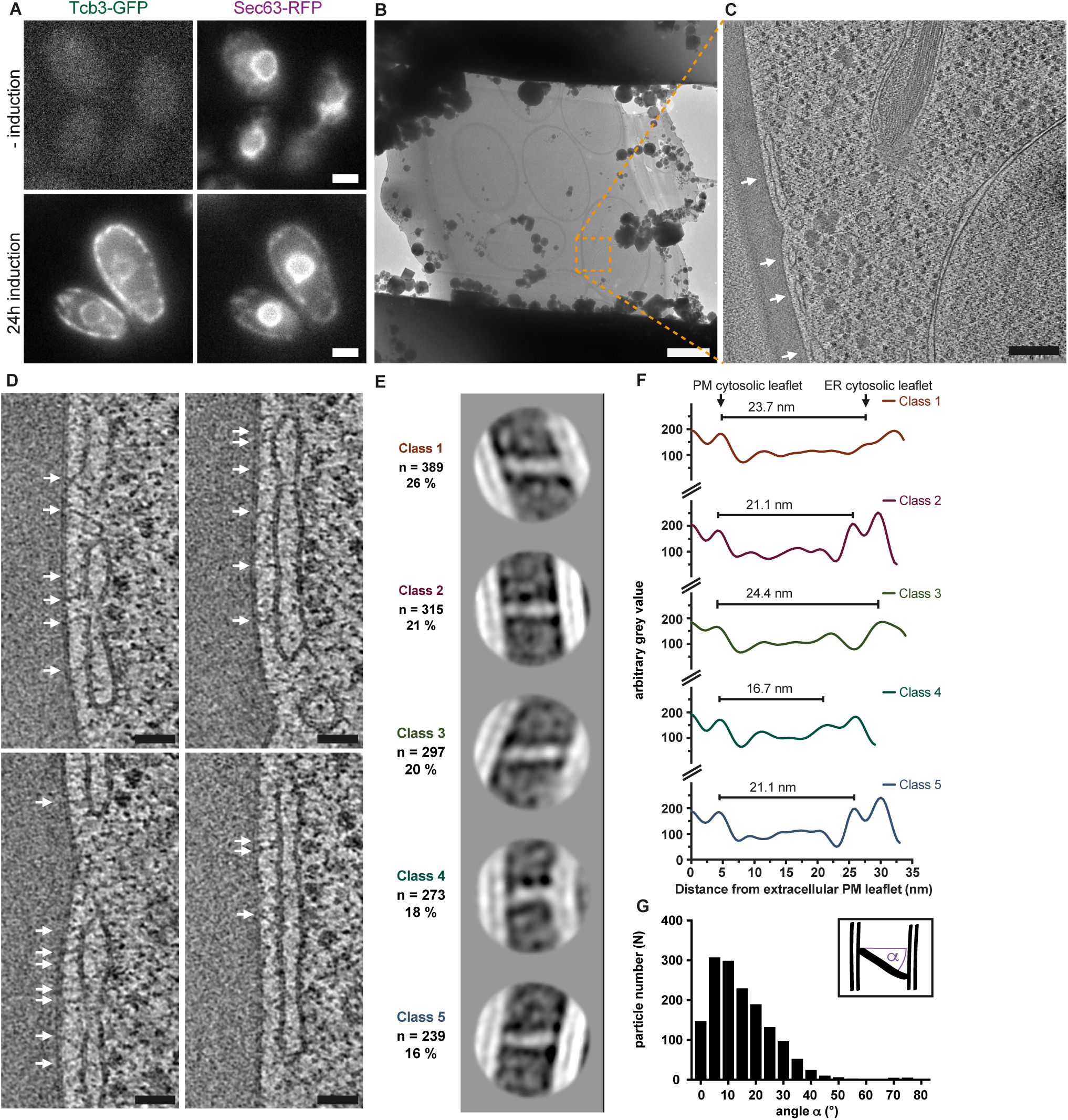
*In situ* structural analysis of Tcb3. **A:** Live FM of *scs2/22*Δ *ist2*Δ *tcb1/2*Δ *prGAL1::Tcb3::EGFP* (5Δ) cells expressing plasmid-encoded Sec63-RFP. Induction of Tcb3-GFP expression rescues ER-PM contact sites, indicated by localization of Sec63-RFP to the cell cortex. **B:** cryo-EM overview of FIB-milled lamella through 5Δ cells after induction. Orange dashed square indicates area corresponding to C. **C:** Virtual slice through electron cryo-tomogram acquired at area indicated in B. Arrows indicate ER-PM contact sites. **D:** Magnified views of virtual slices through electron cryo-tomogram shown in C. Arrows indicate densities bridging ER and PM. **E:** 2D class averages of subvolumes containing bridging particles, grouped into 5 classes. Number of particles per class (n), as well as percentage of total particle set, are indicated. **F:** Projection profiles along major axes of rod-like densities from subtomogram averages. Length measurement indicated for each profile. **G:** Orientation of particle densities vs. particle count; angular deviation α from perpendicular, as depicted in schematic. Scale bars: 2 *µ*m in A and B, 200 nm in C, 50 nm in D.

Cryo-ET of these cells revealed more extended regions of cER as compared to the *scs2/22*Δ *ist2*Δ cells that expressed all tricalbins at endogenous levels (Figure 6B, C). Occasionally, the cER membrane was buckled towards the PM, although the intermembrane distance at these locations was larger than in the *scs2/22*Δ *ist2*Δ cells (5Δ cells: average 16.3 nm, SD = 1.2 nm, N = 10 buckles from 17 tomograms, Supplementary Figure S3). Densities forming bridges between ER and PM were distributed over all the cER (Figure 6D, white arrows). Based on the enrichment of these densities in the 5Δ cells, we reasoned that a large majority of them corresponded to Tcb3-GFP protein particles. While the particles appeared similar in length and thickness, their orientation relative to the plane of the PM varied. The majority of particles deviated from perpendicular by 5-40° (average 15.2°, SD = 12.1°, N = 1513 particles from 17 tomograms) (Figure 6G). We extracted subvolumes containing the particles, collapsed them into a 2D image, and subjected them to subtomogram averaging (see Materials and Methods)(Bharat and Scheres, 2016). Classification resulted in five major, similarly populated classes that together encompassed the vast majority of particles (Figure 6E). The 2D class averages all showed a similar, rod-shaped structure that connected the two membranes nearly perpendicularly (classes 2, 4 and 5), or offset by an angle of 25-30° (classes 1 and 3). In class averages 1-3 and 5, the structures displayed three sequentially arranged major densities (Figure 6E and Supplementary Figure S4). The density closest to the PM appeared tilted relative to the main axis of the particle in these classes. Whereas the densities corresponding to the two leaflets of the PM were similar in all classes, the cER membrane appeared ruffled to various degrees, and displayed a curvature of 34 nm radius in class 4. We estimated the length of the structures by measuring the distance between the cytosolic leaflets of PM and ER using line profiles projected along the structure axes (Figure 6F). In four of the five classes, the distances were 21-24 nm, whereas in class 4 the distance was 16.7 nm. Thus, 82% of the particles were very similar in size and structure, and differed mainly in their orientation relative to the membranes. A small fraction of particles (18%, class 4), were shorter and associated with higher cER curvature. In summary, our analysis suggests that Tcb3 forms a rod-shaped structure that bridges the intermembrane space through three roughly linearly arranged densities. These structures are tilted relative to the PM plane and have distinct lengths.

## Discussion

### ER-PM contact sites are organised into compositionally and ultrastructurally different regions

Mammalian ER-PM contacts mediated by E-Syts appear ultrastructurally distinct from those driving store-operated Ca^2+^ entry, marked by STIM1, suggesting a functional aspect to MCS architecture (Fernández-Busnadiego et al., 2015). Yet, it remains unclear to what extent different MCS proteins segregate within MCS and what the mechanism driving segregation might be. We show here that despite a large degree of coincident localization along cER, different bridging proteins are not distributed homogenously, suggesting that ER-PM contact sites are organised into subdomains of varied composition.

One potential segregation mechanism is that intermembrane distances could drive exclusion of specific proteins based on size (Hoffmann and Kukulski, 2017), as described for cell-cell interactions in T-cell responses and between giant unilamellar vesicles *in vitro* (James and Vale, 2012; Schmid et al., 2016). Distances between the ER and PM are known to be highly variable (West et al., 2011), and it has been hypothesised that different bridging proteins would mediate different intermembrane distances (Gatta and Levine, 2016; West et al., 2011). However, we find that intermembrane distances do not depend on the bridging protein. The average distance was very similar for all bridging proteins. The bridging proteins are thus not the major regulators of intermembrane distance, although they might set constraints to the observed variability. As this variability was similarly large for all bridging proteins, segregation is unlikely to be driven by size-based exclusion.

Instead, our results reveal that the distribution of different bridging proteins correlates with differences in ER curvature. In wild type cells each bridging protein family had a distinct preference for a particular membrane curvature; VAPs and Ist2 localized to cER sheets, whereas tricalbins localized to tubular cER and the curved edges of cER sheets. We suggest that local cER membrane curvature, rather than intermembrane distance, determines bridging protein organisation at ER-PM contact sites (Figure 7A). Localization to cER domains of specific curvature could ensure that the bridging proteins are positioned in proximity to functional partners with similar curvature preference.

**Figure 7.**
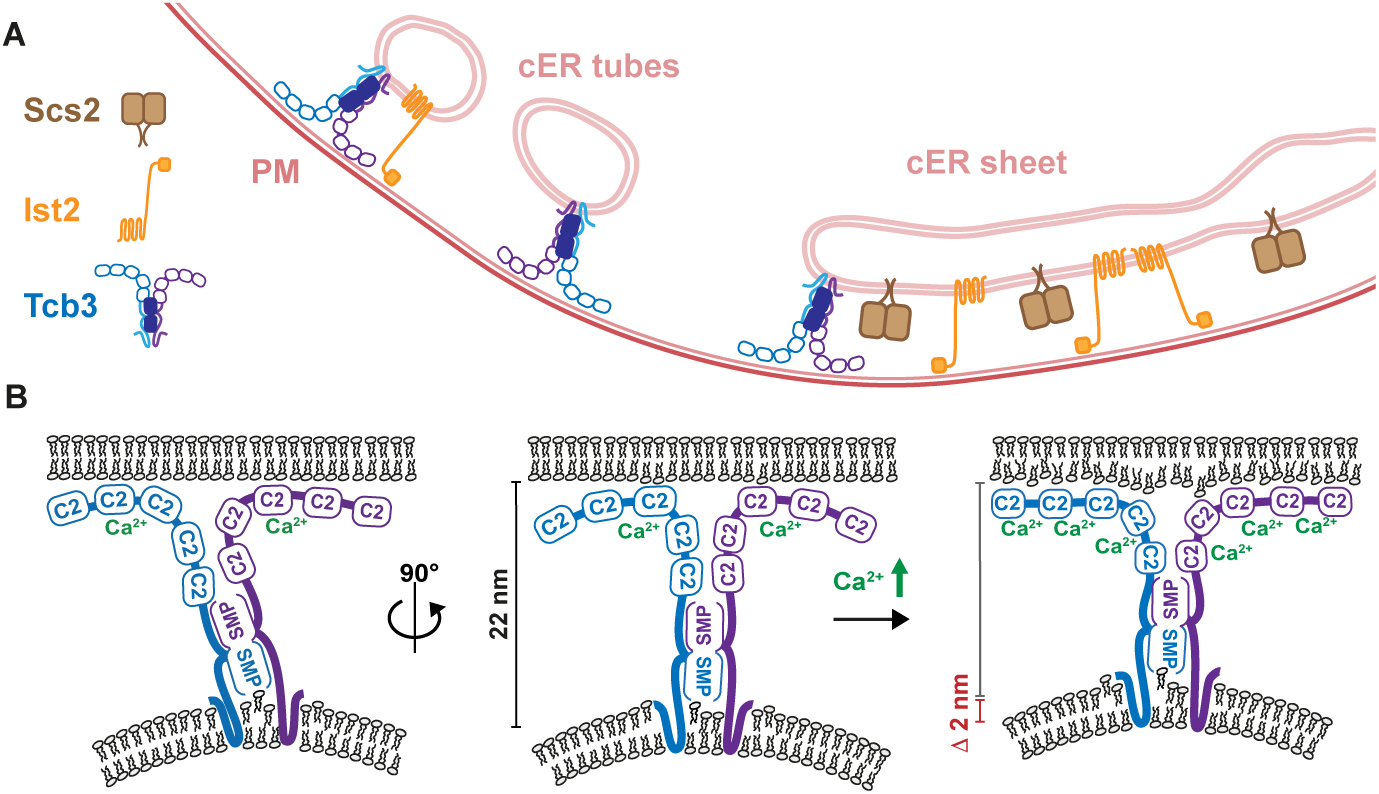
Model for ER-PM contact site architecture and tricalbin organisation. **A:** The distribution of ER-PM proteins correlates with cER membrane curvature, but not intermembrane distance. Tricalbins preferably localize to high cER curvature and are implicated in maintenance of PM lipid asymmetry. **B:** Tcb3 forms a rod-like bridge between the two membranes, and is tilted relative to the PM plane. Its transmembrane domain mediates curvature preference, possibly facilitating lipid extraction/insertion. At a rotated viewing angle, Tcb3 appears perpendicular to the PM plane. It mediates 22 nm intermembrane distance. The dimeric SMP tube arranges vertically to the PM. Not all C2 domains are bound to the PM. Upon increased Ca^2+^ concentrations, more C2 domains bind to the PM, forming a dense membrane coat and potentially introducing bilayer disorder. The intermembrane distance shortens by 2 nm upon increased Ca^2+^.

### Tricalbin-mediated curvature of the cER

The preference of Tcb3 for tubular cER depends on its transmembrane domain, which is predicted to form a hairpin-like structure in the ER membrane similar to the curvature-generating reticulons (Voeltz et al., 2006). In wild type cells Tcb3 overlapped extensively with the pan-ER translocon component Sec63, whereas in the absence of reticulons, Tcb3 and Sec63 segregated into two separate cER pools: sheet-like cER that contained Sec63, and highly tubular Tcb3-positive membranes. Reticulons might modulate overall cER curvature to jointly accommodate curvature-preferring proteins such as tricalbins and bulkier components such as the translocon. Without this modulation, tricalbins either enrich in domains of residual curvature, or generate this curvature themselves. Lipid transfer between membranes *in vitro* is facilitated by high membrane curvature, likely due to bilayer packing defects promoting extraction of lipid molecules (Moser von Filseck et al., 2015)1. The curvature-sensitivity of tricalbins might thus promote lipid transfer between ER and PM (Figure 7B).

### Tricalbins participate in maintenance of PM bilayer asymmetry

Tricalbins and E-Syts are not essential for cell fitness (Manford et al., 2012; Saheki et al., 2016; Toulmay and Prinz, 2012) or mouse physiology (Sclip et al., 2016; Tremblay and Moss, 2016), and even in yeast the lack of phenotype upon deletion has made it difficult to assess their cellular function. We hypothesised that compensatory pathways or redundant protein functions mask phenotypes. Systematic screens using yeast deletion libraries can reveal genetic interactions that provide hints at functional relationships (Costanzo et al., 2016). This technology allowed us to uncover cellular pathways that are specifically dependent on Tcb3, and to assess the importance of the curvature sensing and lipid transfer domains. We found synthetic lethality associated with simultaneous loss of all three tricalbins and other MCS components, supporting the model that MCS function as a cross-regulated network (Lang et al., 2015). Through LTPs that localize to distinct MCS, the intracellular itineraries of lipid molecules are thought to be redundant and can be achieved by multiple routes (Kumar et al., 2018; Lang et al., 2015). Our finding that lack of tricalbins and either of the nuclear-vacuolar junction components Nvj1 or Vac8 leads to reduced growth indicates a redundancy that cannot be compensated further. Another role for tricalbins in lipid flow is reflected in the autophagy-related genes that interacted with tricalbins. Atg2 acts as a tether between the pre-autophagosomal structure (PAS) and the ER to promote expansion of the isolation membrane (Kotani et al., 2018). E-Syts have also been proposed to contribute to autophagosome formation by increasing PI3P synthesis (Nascimbeni et al., 2017). Our finding that autophagy genes, especially Atg2, genetically interact with tricalbins corroborates this model and suggests that ER-PAS contacts involve lipid transfer to expand the forming phagophore.

Of particular interest is the previously unknown role for tricalbins in PM lipid asymmetry, revealed by genetic interactions with RIM101 pathway components. This pathway senses and induces adaption to changes in pH and in the asymmetric distribution of lipids across the PM bilayer, possibly by detecting changes in surface charge (Obara et al., 2012; Rockenfeller et al., 2018). The Rim21 sensory complex assembles at the PM in response to changes in PM asymmetry, triggering the calpain-like Rim13 protease to activate the transcriptional regulator, Rim101 (Obara and Kihara, 2014; 2016). This response ultimately induces expression of PM floppases that re-establish lipid asymmetry in the PM (Ikeda et al., 2008). Although spatially segregated from ER-PM contact sites, the Rim101 pathway is constitutively active in yeast cells lacking all cER (Obara and Kihara, 2016). Our data indicates that this activity is likely due to loss of tricalbins, rather than lack of cER, as our query strains retain ER-PM contacts mediated by Ist2 and VAPs. We propose that tricalbin-mediated lipid transfer contributes to PM asymmetry (Figure 7A), in the absence of which lipid asymmetry is restored through activation of the Rim101 rescue program. The distribution of lipids across the two bilayer leaflets is fundamental to multiple aspects of membrane function, including signalling, membrane potential and curvature generation (Gurtovenko and Vattulainen, 2008; McMahon and Boucrot, 2015; Segawa and Nagata, 2015). Precisely which lipids are affected by loss of tricalbin function remains to be determined, and will require novel methods for dissecting specific lipid localization.

### Tcb3 curvature-generating and lipid-transfer domains are functionally dispensable

Surprisingly, Tcb3 function could largely be rescued by mutants in which we replaced its curvature-sensitive or its lipid transfer domain. We consider that either these molecular functions are not relevant for cellular roles of Tcb3, or the two domains perform redundant functions. Indeed, reticulons drive ER membrane curvature and facilitate lipid transfer (Voss et al., 2012), and thus might compensate when Tcb3 loses its curvature-sensitive properties. Similarly, lipid transfer via the SMP domain (Saheki et al., 2016) may be replaced by other LTPs with related lipid binding domains, such as Vps13 (Kumar et al., 2018). Finally, the two domains of Tcb3 might be functionally redundant with each other: effects on lipids at Tcb3-mediated contacts might be facilitated either by curvature or by SMP function. In light of the genetic interactions we uncovered, further focus on the C2 domains of Tcb3 is a priority. Among ER-PM bridging proteins these Ca^2+^-responsive domains are unique to the tricalbins and thus are likely to be functionally important.

### C2 domain organisation at the PM is Ca^2+^ dependent

In support of an important role for tricalbin C2 domains, Ca^2+^ stimulation had marked effects on tricalbin-mediated contacts, resulting in increased density on the PM, reminiscent of a protein coat. The thickness of this putative coat is compatible with a layer of densely packed C2 domains. Previous biochemical data suggested that Ca^2+^ binding displaces a E-Syt1 C2 domain from an occluding position on the SMP channel, relocating it to the PM (Bian et al., 2018). Many C2 domains bind PIP2 in a Ca^2+^-dependent manner, thereby regulating interaction with target membranes (Schiavo et al., 1997). Our data provide *in situ* evidence supporting that Ca^2+^ enhances binding of tricalbin C2 domains to the PM. Interactions of C2 domains with lipids can change bilayer structure as well as induce membrane curvature (Martens et al., 2007). Thus, engagement of the tricalbin C2 domains with the PM could facilitate insertion or extraction of lipid molecules by inducing disorder in the bilayer (Figure 7B). A second effect of high Ca^2+^ was a shortening of the distance between ER and PM from 23 to 21 nm. This is a more modest change than has been observed in mammalian cells upon thapsigargin treatment, which induced a narrowing of E-Syt1 mediated contacts from 22 to 15 nm (Fernández-Busnadiego et al., 2015). Together, our data suggest that under normal Ca^2+^ conditions, not all C2 domains of the tricalbins are bound to the PM. At high Ca^2+^, coincident with reduced ER-PM distance, more C2 domains bind to the PM, and at maximal binding form a densely packed coat (Figure 7B).

### Tcb3 forms a rod-shaped structure with linear domain arrangement

Subtomogram averaging of densities from contacts mediated by Tcb3 alone permits a detailed *in situ* structural analysis, suggesting that Tcb3 forms remarkably rigid, rod-like structures that are mostly oriented non-perpendicular to the PM plane. Different 2D class averages shared significant similarities, suggesting that they mostly represent different views of a single structure. This would imply that Tcb3 is tilted by approximately 20° at its highest inclination, appearing perpendicular at views rotated by 90° round the membrane normal (Figure 7B). The lengths of the average particles in the respective classes are consistent with this model.

Subdivision of average structures suggests that the cytosolic domains of Tcb3 are arranged in a linear fashion along the axis of its rod-like structure (Figure 7B). These domains include the SMP domain, which forms a barrel about 4.5 nm long and 3 nm wide. Previous structural data show that SMP domains form head-to-head dimers resulting in 9 nm long tubes, indicating that dimerization is a general feature of SMP domain organisation (AhYoung et al., 2015; Kawano et al., 2018; Schauder et al., 2014). The central densities in our particle averages are compatible with an SMP dimer orientated along the particle axis (Figure 7B). This arrangement excludes a model in which the SMP dimer forms a shuttle parallel to the membrane. This model has been favoured because a central density parallel to the membranes was observed in E-Syt1 overexpression conditions, and was enhanced upon Ca^2+^ stimulation (Fernández-Busnadiego et al., 2015; Reinisch and De Camilli, 2016). Although we also detect extra density at high Ca^2+^, the layer we observe lies immediately adjacent to the PM. This coat-like density is similar to PM-bound BAR domains (Bharat et al., 2018), and is suggestive of an intimate interaction with the PM, likely to be driven by Ca^2+^-responsive C2 domains rather than SMP domains.

An open question about lipid transfer at MCS is the mode of lipid travel across the soluble interface between bilayers. One model is direct tunnelling of lipid molecules from one to the other bilayer through the 9 nm SMP dimer barrel, which requires intermembrane distances in the range of 10 nm (Schauder et al., 2014). We measured the shortest intermembrane distances at positions of local cER buckling towards the PM to be ∼14 nm, which was similar to the length of the shortest structure in 2D class averages. Over larger contact areas, the measured distances of 21-24 nm corresponded to the length of most of the particles. Thus, under all circumstances we analysed, the distance across which lipid molecules must be transferred was too large to be compatible with tunnelling via SMP dimers alone. This could mean that tunnelling is a transient event that we fail to capture in our analysis. Alternatively, an additional transfer mechanism could bridge the remaining 5-12 nm.

The unprecedented *in situ* structural view on tricalbins bridging two organelles provides an essential basis for addressing the molecular mechanism of lipid transfer. Our CLEM results reveal how specific the ultrastructure of ER-PM contacts is depending on the bridging proteins, implying that contact site function is regulated through variability in composition as well as architecture, potentially adjusted to their cellular roles. We uncover such a role for tricalbins in maintenance of bilayer asymmetry in the PM. Our study shows a way to untangle the intricate network of redundancies at the protein domain and pathway level, which emerge as a fundamental principle of the cell’s overlapping lipid routes.

## Acknowledgements

We thank the Electron Microscopy and Light Microscopy Facilities of the MRC-LMB for support in data collection, Jake Grimmett and Toby Darling for computing support, Julia Mahamid for instructions on cryo-FIB milling, Christopher Russo for help in using the Scios microscope, Jerome Boulanger for the script to analyse intermembrane distances. We thank Ruben Fernandez-Busnadiego and Javier Collado for sharing unpublished results. We thank Pablo S. Aguilar for the kind gift of the plasmid encoding GCamP, Susan Michaelis for the plasmid encoding Sec63-RFP, Wendell Lim and Kurt Thorn for the plasmid encoding mRuby2. This work was supported by the Medical Research Council (MC_UP_1201/8 to W.K. and MC_UP_1201/10 to E.A.M.). T.A.M.B. is funded by a Sir Henry Dale Fellowship from the Wellcome Trust and the Royal Society (Grant No. 202231/Z/16/Z) and a Vallee Foundation Scholarship.

## Materials and Methods

### Yeast strains

Chemically-competent yeast cells were prepared from cultures grown in standard YPD medium at 30°C. Cultures of strains for fluorescent live imaging or EM sample preparation were inoculated from overnight pre-cultures and grown to mid-log phase at 25°C in 10 - 50 ml strain-specific selective dropout medium without tryptophan. All cells were grown with 2% glucose as carbon source with exception of WKY308, the galactose-inducible Tcb3-EGFP 5Δ strain, which was cultivated in synthetic complete media lacking tryptophan (SC-Trp) with 1% raffinose and induced by addition of 1% galactose.

### Yeast genetic techniques

Yeast strains (see Table S1) were generated by homologous recombination of the target genes with PCR cassettes for gene deletion or tagging as described in (Janke et al., 2004). Seamless N-terminal tagging with sfGFP was performed using the pMaM173 plasmid and yeast strains with Gal L-ISCE1 integration to loop out the URA marker as described in (Khmelinskii et al., 2011). Individual transformants were isolated on selection plates and insertion of PCR cassettes was verified by colony PCR. Deletion strains were generated by homologous recombination of target genes with PCR deletion cassettes from plasmids pFA6a-natNT2, pFA6a-hphNT1, pFA6a-klURA3, pFA6a-Leu2 and pFA6a-KanMX6. Yeast strains carrying multiple deletions were generated by either subsequent homologous recombination or by mating and tetrad dissection. A galactose-inducible Tcb3-EGFP strain was generated by insertion of the KanMX6-GAL1pr PCR cassette amplified from pFA6a-KanMX6-GAL1pr, which introduced the GAL1 promoter 5’ of the endogenous *TCB3* locus. By mating and tetrad dissection, a galactose-inducible Tcb3-EGFP strain deleted for *tcb1/2 scs2/22 ist2* was isolated. A GCamP reporter strain deleted for *scs2/22 ist2* was generated by homologous recombination of the PvuII linearized PSAB367 plasmid (Carbó et al., 2017) at the *URA3* locus and selected for on G418 selection plates. All yeast transformations were performed using a standard lithium acetate transformation protocol.

### Constructs and Cloning

The *tcb3* gene including 500bp of the 5’ UTR and 100 bp of the 3’ UTR was amplified from purified genomic DNA using Phusion DNA polymerase (New England Biolabs). 5’Sal1 and 3’Sac1 restriction sites were introduced with amplification primers. Amplified DNA was ligated with linearized pRS316 plasmid at Sal1 and Sac1 restriction sites. A C-terminal EGFP cassette was introduced by homologous recombination in yeast of the amplified EGFP sequence with homology arms to the 3’ end of *tcb3* and 3’ UTR to the linearized pRS316-Tcb3-URA3 vector. Chimera-Tcb3-EGFP was generated by homologous recombination of the Afe1 linearized pRS316-Tcb3-EGFP-URA3 and PCR-amplified sequence of *erg11* TMD domain (1-55 aa-3x-gly) with homology arms directly targeting the regions before the start codon and 742 bp within the sequence of *tcb3.* This resulted in replacement of the N-terminal 247 aa of Tcb3. The SMP domain (272-479 aa) of Tcb3 was replaced by homologous recombination with a synthetic DNA primer with (GGGGS)_2_ linker sequence and homology arms targeting 813 bp and 1438 bp within *tcb3* sequence. For C-terminal tagging with mRuby2, we removed the 5’ linker of the mRuby2 sequence from pFA6a-link-yomRuby2-KanMX (Lee et al., 2013). A Sal1 restriction site was introduced directly 5’ of the mRuby2 sequence which removed 29 bp to the S3 primer binding site. Following selection, plasmids were recovered from construct-bearing yeast strains, by cell lysis with glass-beads and plasmid prep with the QIAprep Spin Miniprep Kit. Plasmids were re-transformed into calcium-competent bacteria for amplification. Correct insertion and sequence for all inserts was validated by DNA sequencing (GATC Biotech).

### Live-cell imaging

Yeast cells were grown to mid-log phase in SC-Trp or strain-specific selective dropout medium lacking tryptophan at 25°C. Cells were adhered to glass coverslips using ConcanavalinA and imaged in SC-Trp medium using a Nikon TE2000-E widefield microscope, which was controlled by Nikon NIS Elements 4.4 and equipped with 100x oil-immersion TIRF objective with NA 1.49, NEO sCMOS DC-152Q-C00-FI camera (Andor), Lambda DG-4 lamp (Sutter Instruments), filter sets 49002 ET GFP(Chroma), excitation 470/40, dichroic T495lpxr, emission 525/50 for GFP, 49005 ET DSRED (Chroma), excitation 545/30, dichroic T570Ipxr, emission 620/60 with additional emission filter 605/70 for RFP, 49006 ET CY5 (Chroma), excitation 520/60, dichroic T660Ipxr, emission 700/75 for far red. Images were acquired with 0.5-3 s exposure time, depending on fluorescence intensity of target proteins. Representative images used in Figure panels were adjusted individually for best contrast and brightness in Fiji and therefore do not represent quantitative differences in fluorescence intensity, with exception of Supplementary Figure S2C, which was adjusted for quantitative comparison. Line profiles to visualize colocalization were extracted around the cell cortex and straightened using Fiji.

### SGA analysis

Synthetic Genetic Array (SGA) analysis was performed according to (Tong and Boone, 2007) using a colony pinning robot (Singer Instruments). The *tcb1/2*Δ query strain (WKY330) or *tcb1/2/3*Δ query strain (WKY331) without or with Tcb3 rescue constructs were mated to the haploid deletion mutant array and Decreased Abundance by mRNA Perturbation (DAmP) array at a density of 384 spots per plate. Diploid strains were selected, then pinned onto sporulation medium and incubated for 5-7 days at 25°C. Haploid *MATa*, triple mutants for the *tcb1/2*Δ query strain, quadruple mutants for the *tcb1/2/3*Δ query strain, and rescue mutant strains were selected on appropriate selection media. Plate supplements were used at the following concentration: 50 mg/L Canavanine, 50 mg/L S-Aminoethyl-L-cysteine, 200 mg/L G418, 100 mg/L ClonNAT, 300 mg/L HygromycinB. Synthetic sick and lethal phenotypes were identified using PhenoBooth plate imager (Singer Instruments) and PhenoSuite colony counting and analysis software (Singer Instruments). Colony growth on selection plates was scored and compared to colony growth on *MATa* control plates. Colony size was normalized per plate and row and a log2 ratio for selected deletion mutants was calculated. Reduction of growth by factor of 2 (log2 ratio less than −1) was used as a cut-off for synthetic sick phenotypes. To verify selected hits, gene deletion mutants in the *tcb1/2/3*Δ background and single deletion mutants were isolated by tetrad dissection or generated anew by homologous recombination with gene deletion cassettes and subsequent verification. For serial dilution growth assays, late log-phase cultures were diluted to OD^600^ of 2 before preparing 10-fold serial dilutions. A volume of 3 µl per dilution was spotted on plates as indicated in Figure 4. Because Rim101 pathway mutants are sensitive to growth under high salt conditions (Hayashi et al., 2005), growth of these mutants was assayed on YPD with 0.1 M LiCl.

### Room-temperature correlative light and electron microscopy

Room-temperature CLEM was performed as described in (Kukulski et al., 2011) with modifications described in (Ader and Kukulski, 2017). Yeast cells were grown to logarithmic phase in SC-Trp medium as described before and pelleted by vacuum filtration. Resulting yeast paste was high-pressure frozen in 200 µm deep wells of aluminium carriers (Wohlwend) using a HPM100 (Leica Microsystems). Freeze-substitution and Lowicryl HM20 (Polysciences, Inc.) resin-embedding followed the protocol and timing described in (Kukulski et al., 2011), except that 0.03-0.05% uranyl acetate in acetone was used for freeze-substitution. During some preparations, samples were shaken on dry ice for the first 2 h of freeze-substitution. Sections of 270 nm thickness were cut with an Ultra 45° diamond knife (Diatome) on an Ultracut E microtome (Reichert). The sections were floated onto PBS and picked up with 200 mesh carbon-coated copper grids (AGS160, Agar Scientific). Fluorescent TetraSpeck (Invitrogen) beads, 50 nm in diameter, were absorbed onto the grid. Directly after sectioning, grids were mounted for fluorescence microscopy in PBS and imaged as previously described for live-cell imaging. Prior to ET, 15 nm gold beads were adsorbed on the sections, which were then post-stained for 10 min with lead citrate. Scanning transmission EM tomography was done on a TF20 microscope (FEI) with an axial brightfield detector, using a camera length of 200 mm and a 50 µm C2 aperture (Ader and Kukulski, 2017; Hohmann-Marriott et al., 2009). For low magnification correlation either montages or tilt series at 4.4 nm pixel size were acquired using SerialEM (+/- 60° tilt range, 2° increment, single axis acquisition) (Mastronarde, 2005). Higher magnification dual axis tilt series were acquired with 1° increment and at 1.1 nm pixel size (Mastronarde, 1997). All tomographic reconstructions were done in IMOD (Kremer et al., 1996) and fiducial-based correlation was done using MATLAB-based scripts described in (Kukulski et al., 2011).

### Classification and statistical analysis of ER-PM contacts identified by RT-CLEM

In tomograms of resin-embedded cells, ER morphologies at ER-PM contacts associated with signals of the different GFP-tagged proteins were visually classified into four different classes. The first class contained fluorescent signals spread over ER sheets or cisternae (“flat ER sheet”). In the second class, fluorescent signals localized to ER tubes (“tubular ER”). Isolated punctate fluorescent signals localizing to curved ends of ER sheets or fenestrations were also included in this class. Thirdly, ER of more complex morphology, which contained both tubular and flat parts throughout the volume, were counted as a third class (“mixed flat & tubular”). Fluorescent signals that could not be assigned to any ER membrane were classified as “n.a.”. To show the 3D membrane morphology in the Figure panels, segmentations of the cER membrane, along the cytosolic leaflet, throughout the tomographic volume were done in Amira (Thermo Fisher). For better visualization of the cER morphology, the segmentation models in Figures are displayed tilted and rotated with respect to the orthogonal view of the tomogram. In total, the RT-CLEM dataset on sfGFP-Scs2 cells contained 42 correlated fluorescent signals in 15 cells, on sfGFP-Ist2 cells 45 correlated signals in 14 cells, on Tcb3-EGFP cells 55 correlated signals in 18 cells, on Tcb3-EGFP *scs2/22*Δ *ist2*Δ cells 58 correlated signals in 25 cells, on sfGFP-Ist2 *tcb1/2/3*Δ *scs2/22*Δ cells 69 correlated signals in 25 cells, on Tcb3-EGFP *rtn1*Δ *yop1*Δ cells 40 correlated signals in 12 cells and 8 areas devoid of Tcb3-EGFP signals at sheets in 3 cells. Resin sections from different freeze-substitution and embedding runs varied in preservation quality, which resulted in variances in fluorescence and visibility of membrane ultrastructure. For this reason, correlations to areas with poor preservation of membrane ultrastructure preservation were excluded from intermembrane distance measurements. Intermembrane distances at ER-PM contacts were measured approximately 100 nm around the extent of fluorescent signals, according to the transformation based on fiducial markers. Points along the cytosolic leaflets of the ER membrane and of the PM were clicked in IMOD were the membrane was best visible, approx. every 10 nm in the x/y plane. The membranes were traced every 5.5 nm in z direction throughout the tomographic volume. Interpolated surfaces were fit through the sets of points in MATLAB. The minimum and maximum distance, the average distance and standard deviation were calculated between the PM surface model and the sets of points clicked at the ER membrane. The output of the surface model was plotted and visually inspected using MATLAB. The average membrane distance of each ER-PM contact associated with a fluorescent signal was treated as one individual observation (N) and the average and the standard deviation (SD) of all observations for each analyzed yeast strain were combined and compared.

### Cryo-EM sample preparation

Yeast cells were grown to mid-log phase in SC-Trp medium. One OD_600_ unit of cells was pelleted by centrifugation at 3000x g. Cells were resuspended in 500 µl SC-Trp containing 15% high-molecular weight dextran (w/v) and 2% glucose or 1% raffinose for cells with induced protein expression. Vitrification by plunge freezing was done with a manual plunger and a temperature controlled liquid ethane reservoir at −181°C (Russo et al. 2016). 5 µl cell suspension were applied to Quantifoil R2/2 Cu 200 mesh or UltrAuFoil R2/2 Au 200 mesh, which beforehand were plasma treated for 40 sec using Fischione 1070 Nano Clean with 9/1 argon/oxygen mix and 38 W power. Grids were backside-blotted for 12-15 s with filter paper grade1 (Whatman) immediately before plunge freezing.

### Cryo-FIB milling

Cryo-FIB milling of plunge frozen yeast cells was performed on a Scios DualBeam FIB/SEM microscope (FEI) equipped with a Quorum PP3010T cryo-FIB/SEM preparation system. The loading stage and milling procedure were adapted, with minor alterations, from (Schaffer et al., 2015). The temperature of the cryo-stage was kept between −170°C to −180°C and the anti-contaminator temperatures below - 190°C. Grids were sputter-coated with platinum in the Quorum preparation system for 60 s at 10 mA current before milling. Additionally, grids were coated with a layer of organometallic platinum using the gas injection system for 30 s at a distance of 5 mm with a stage tilt of 25°. Imaging settings for the electron beam were 2kV voltage and 13 pA current and for the ion beam 30 kV voltage and 10 pA current. Initial rough milling was performed with 30 kV ion beam voltage and 0.5 or 1 nA current at a stage tilt of 35°-40°, below and above the cells of interest. The stage was then tilted to 17° and cells were subsequently milled to 3 µm lamella thickness with 30 kV, 0.3 nA, then to 1 µm lamella thickness with 0.1 nA. Usually 2-3 lamellae were rough-milled per grid before proceeding to a fine milling step. Fine milling was performed by milling either at 16° or 18° stage tilt as described by (Schaffer et al., 2017) until the lamella was approx. 300 nm thick. Final polishing of the lamellae was performed at 17° stage tilt and resulted in 150-300 nm thick lamellae with 10° pre-tilt relative to the grid. All fine milling steps were done at 16 kV voltage and 11 pA or 23 pA ion beam current to minimize potential ion beam damage to the lamellae.

### Cryo-fluorescence microscopy

To target yeast cells with high levels of intracellular Ca^2+^, before plunge freezing cells were resuspended in 500 µl SC-Trp containing, 15% high molecular weight dextran (w/v), 2% glucose and 200 mM calcium chloride. Cells were plunge-frozen as described above, within a 2-10 min time window after resuspending in the calcium chloride containing medium. Cryo-fluorescence imaging of yeast cells on TEM grids was performed with few adaptation according to (Ader et al., 2019) on the Leica EM cryo-CLEM system with an HCX PL APO 50x cryo-objective with NA = 0.9 (Leica Microsystems), an Orca Flash 4.0 V2 SCMOS camera (Hamamatsu Photonics), a Sola Light Engine (Lumencor) and the L5 filter, excitation 480/40, dichroic 505, emission 527/30 for detection of GFP fluorescence. The room humidity was kept between 20-25% and the microscope stage was cooled to −195 °C during imaging. Using the Leica LAS X software, a 1.5 × 1.5 mm tile scan z-stack with 18 µm range and 2 µm step size was recorded around the center of the grid in the GFP channel (17% intensity, 3 s exposure) and in the brightfield channel (30 intensity, 50 ms exposure). Autofocus routine was performed in the brightfield channel. Groups of cells with the highest fluorescent signals were identified by adjusting the contrast and brightness in Fiji. These groups of brightest cells were targeted for cryo-FIB milling by visual correlation using overview SEM images for identification of cells. Overlays of cryo-FM and cryo-EM images were generated using the eC-CLEM plugin in Icy (Paul-Gilloteaux et al., 2017).

### Electron cryo-tomography

Cryo-ET data acquisition was done on two Titan Krios microscopes (Thermo Fisher) fitted with Quantum energy filter and K2 direct electron detector (Gatan) in counting mode, using SerialEM (Mastronarde, 2005). Montaged images of the central part of the grid were acquired at a pixel size of 200 nm to localize the lamellae on the grid. Overview montages of the individual lamellae were acquired at pixel sizes of 5.4 or 5.5 nm, depending on the microscope, to assess the lamellae quality and identify ER-PM contact sites within the cells. Tilt series were collected between ±60° starting from 0° using a grouped-dose symmetric tilt scheme with 1° increment and group size of 4 (Bharat et al., 2018; Hagen et al., 2016). Cryo-ET datasets were acquired during multiple sessions, during which the calibrated pixel size at which tilt series were acquired varied between 3.54 Å and 3.77 Å. For analysis of all datasets, the average pixel size 3.7 Å was used. The target dose rate was kept at ∼2-4 e^-^/pixel/s on the detector depending on lamella thickness. The energy filter slit width was set to 20 eV. The nominal defocus was varied between −3.5 µm and −6 µm for different tilt series. A dose of approximately 1.0-1.3 e^-^/Å^2^ was applied per image of tilt series. Tilt images were collected as frames, and frames were aligned using the alignframes program in IMOD. Tilt series were aligned using patch tracking and reconstructed using IMOD. The contrast transfer function was estimated and compensated for by phase flipping in IMOD. Tomograms were reconstructed in IMOD by backprojection for image processing (see below), and by simultaneous iterative reconstruction technique (SIRT) with 10 iterations at a pixel size of 7.4 Å for display in Figures. In addition, to improve visibility of low-resolution morphological features, nonlinear anisotropic diffusion filtering (Frangakis and Hegerl, 2001) in conjuction with gaussian filtering was applied to tomographic slices.

### Image processing and classification of cryo-ET data

To analyze the density layer observed at the PM of ER-PM contacts at high cytosolic calcium, a subtomogram averaging approach described previously (Bharat et al., 2018) was adapted and used. First, the 4 (out of 10) tomograms of the high calcium dataset that displayed a visible density layer were rotated around their x axis in IMOD to display the PM perpendicularly to the viewing plane. Next, along the inner leaflet of the PM where a dense coat was visible, points were clicked in slices spaced by 8 nm in z direction. For comparison, the same was done at the PM outside the ER-PM contact site in the same tomograms. A spline was fitted through each set of points in MATLAB and subtomograms were extracted along the fitted spline in overlapping boxes using RELION (Bharat and Scheres, 2016). The in-plane rotation angle from the spline fit was written out and retained for subsequent alignment. The subtomograms were projected into 2D images and subjected to 2D averaging using RELION (Scheres, 2012) separately for each area of PM. Several classifications were performed for each membrane profile with different parameters to control for spurious alignments. Selection of well-aligned classes was based on visual examination of results from different runs and different membrane areas. Classes from PM-ER contacts with the extra layer were compared to free PM for each tomogram individually, to minimize defocus effects on the appearance of the membrane bilayer that vary between tomograms. To measure the extent of the extra layer in the 4 tomograms of cells with high cytosolic calcium and in 5 tomograms of the control cells, the volume of cER and the volume occupied by the extra layer in the tomograms were segmented and calculated in Amira. The ratio of the extra layer volume to the volume of cER was determined and served as a measure of the extent of the extra layer.

Individual particle densities, which appeared to bridge the ER and PM in tomograms of *scs2/22*Δ *ist2*Δ *tcb1/2*Δ cells overexpressing Tcb3-GFP, were manually picked in IMOD by clicking one point at the ER membrane base, a second point at the PM, and two auxiliary points approx. 10 nm away from the bridging density along the ER membrane and along the PM. This set of points allowed calculating the in-plane angle relative to the membranes for each bridging density. Subtomograms were extracted at the center of each density using RELION and the in-plane rotation angle for each subtomogram was written out and retained for the subsequent image alignment. Subtomograms were projected into 2D images and subjected to 2D averaging in RELION. Several classifications with varying parameters including the number of classes were performed and well-aligned classes within the different runs were compared. Varying the parameters of different runs resulted consistently in the similar class averages, of which one representative run with 5 class averages is shown in Figure 6. The dataset for 2D averaging of bridging densities in *scs2/22*Δ *ist2*Δ *tcb1/2*Δ cells overexpressing Tcb3-GFP consisted of 17 electron cryo-tomograms, from which 1513 bridging densities were selected based on their visibility approx. parallel to the viewing plane. Line profiles of class averages were made in Fiji along the major axes of the average particles over a width of 15 pixels, which included the whole width of the average particles.

Average ER-PM intermembrane distances in electron cryo-tomograms were measured in the same manner as described before for the RT-CLEM dataset. At positions where ER membrane buckled towards PM, local distances were measured manually at the shortest distance between the PM and the ER membrane using IMOD.

### Quantification and Statistical Analysis

Quantitative analysis of membrane morphology and membrane distances are described in the Methods details for the individual experiments and contains information about dataset size, mean, SD and N. Statistical significance was tested in using Welch’s unequal variances *t*-test in Graphpad Prism.

---

**Supplemental Figure S1.**
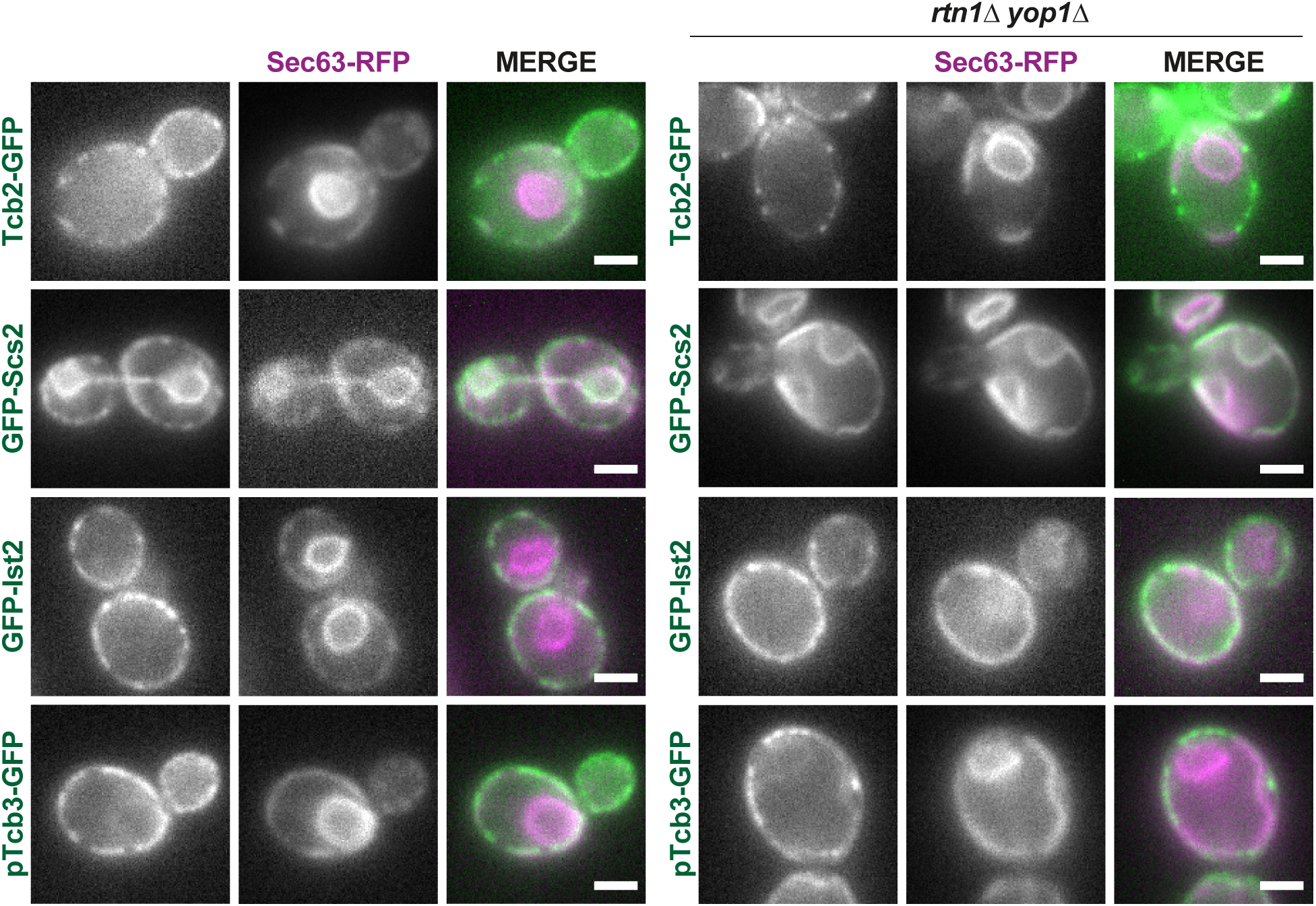
Reticulon-dependent localization of ER-PM proteins. Live FM of wild type (left panel) and *rtn1*Δ *yop1*Δ (right panel) cells expressing either Tcb2-EFGP, GFP-Scs2, GFP-Ist2 or plasmid-encoded Tcb3-GFP, in combination with plasmid-encoded Sec63-RFP. Scale bars: 2 *µ*m.

**Supplemental Figure S2.**
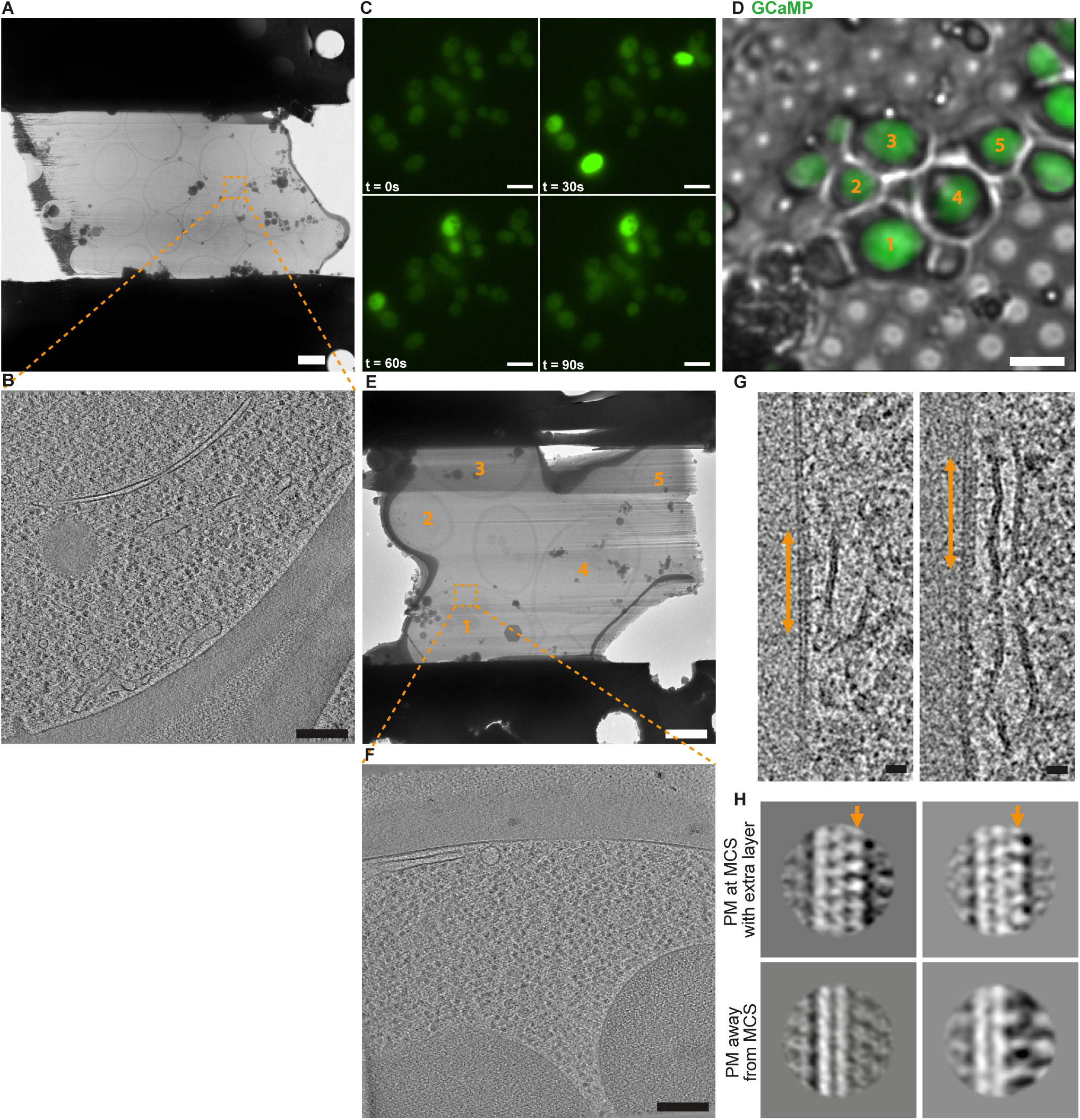
Cryo-CLEM for targeting cells with high cytosolic Ca^2+^ by cryo-FIB milling and cryo-ET. **A:** Overview cryo-EM image of a lamella prepared from untreated *scs2/22*Δ *ist2*Δ cells. Orange dashed square indicates the field of view of the electron cryo-tomogram shown in B. **B:** Orthogonal virtual slice through electron cryo-tomogram acquired at the lamella position indicated in A. Left image in Figure 5A is from the same electron cryo-tomogram. **C:** Live FM of a group of *scs2/22*Δ *ist2*Δ cells expressing GCaMP. GCaMP signal (green) imaged at different time points (0s, 30s, 60s and 90s) after exposure to 200 mM CaCl_2_. **D:** Cryo-FM image of *scs2/22*Δ *ist2*Δ cells expressing GCaMP; overlay of green and bright field signals. Cells that display strong GCaMP signals, and that are visible in the FIB-milled lamella (E) of this cell group are labeled by matching numbers 1 – 5 in both D and E. **E:** Cryo-EM overview of FIB-milled lamella generated from the group of cells shown in D; labeled by matching numbers. Orange dashed square indicates the field of view of the electron cryo-tomogram shown in F. Note that this image corresponds to the image in Figure 5B, but is shown here without overlayed GCaMP signal. **F:** Orthogonal virtual slice through electron cryo-tomogram acquired at the lamella position indicated above in cell no. 1. Magnified views of virtual slices from this tomogram are shown in Figure 5C left and middle. **G:** Virtual slices of electron cryo-tomograms of cells displaying strong GCaMP signals, examples showing an extra density layer (orange arrows) in addition to those shown in Figure 5C. Left panel is a different slice from the same tomogram as shown in middle panel of Supplemental Figure S3A. **H:** 2D class averages from subtomogram averaging of PM within ER-PM contact with extra layer (top images), and PM outside of ER-PM contact (bottom). Class averages to the left and right correspond to tomogram shown in left and right image of G, respectively. In class average images, extracellular space is to the left, cytosol to the right. Scale bars: 2 µm in A and E, 200 nm in B and F, 5 *µ*m in C and D, and 20 nm in G.

**Supplemental Figure S3.**
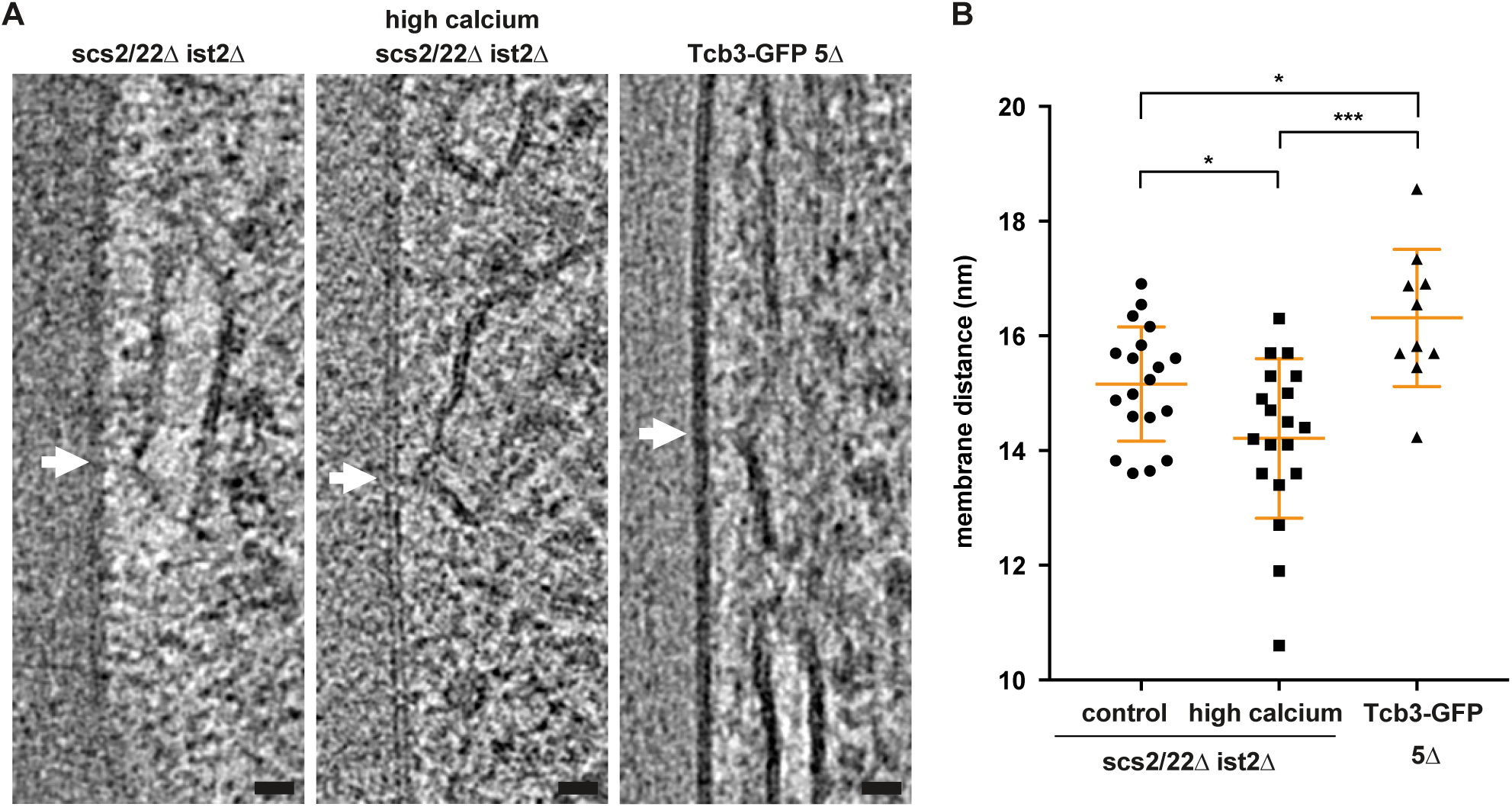
The cER locally buckles towards the plasma membrane. **A:** Virtual slices through electron cryo-tomograms of three different conditions; untreated *scs2/22*Δ *ist2*Δ cells, Ca^2+^-treated *scs2/22*Δ *ist2*Δ cells displaying strong GCaMP signal, and 5Δ cells expressing galactose-induced Tcb3-GFP. Arrows indicate local buckles in cER. Middle panel is a different slice through the same tomogram as shown in left panel of Supplemental Figure S2G. **B:** Distances between the buckling cER and the plasma membrane, measured locally at positions of buckling. Orange lines represent mean and SD. (untreated vs. treated: P=0.0213, untreated vs. 5Δ cells: P=0.0192, treated vs. 5Δ cells: P=0.0004)

**Supplemental Figure S4:**
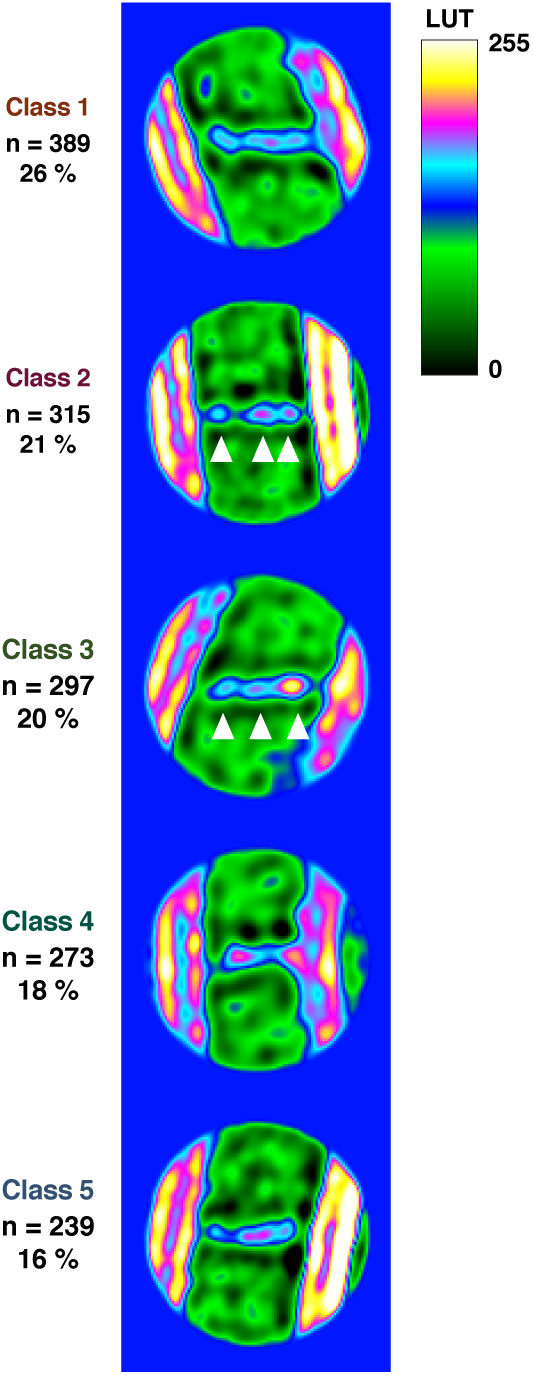
2D class averages reveal linear domain organization of rod-like particles. 2D class averages of subvolumes containing bridging particles as shown in Figure 6E, but grey scale was converted to heat map for better visibility of density variations. LUT indicates color corresponding to gray scale values. Number of particles per class (n), as well as percentage of total particle set, are indicated as in Figure 6E.

**Table S1:**
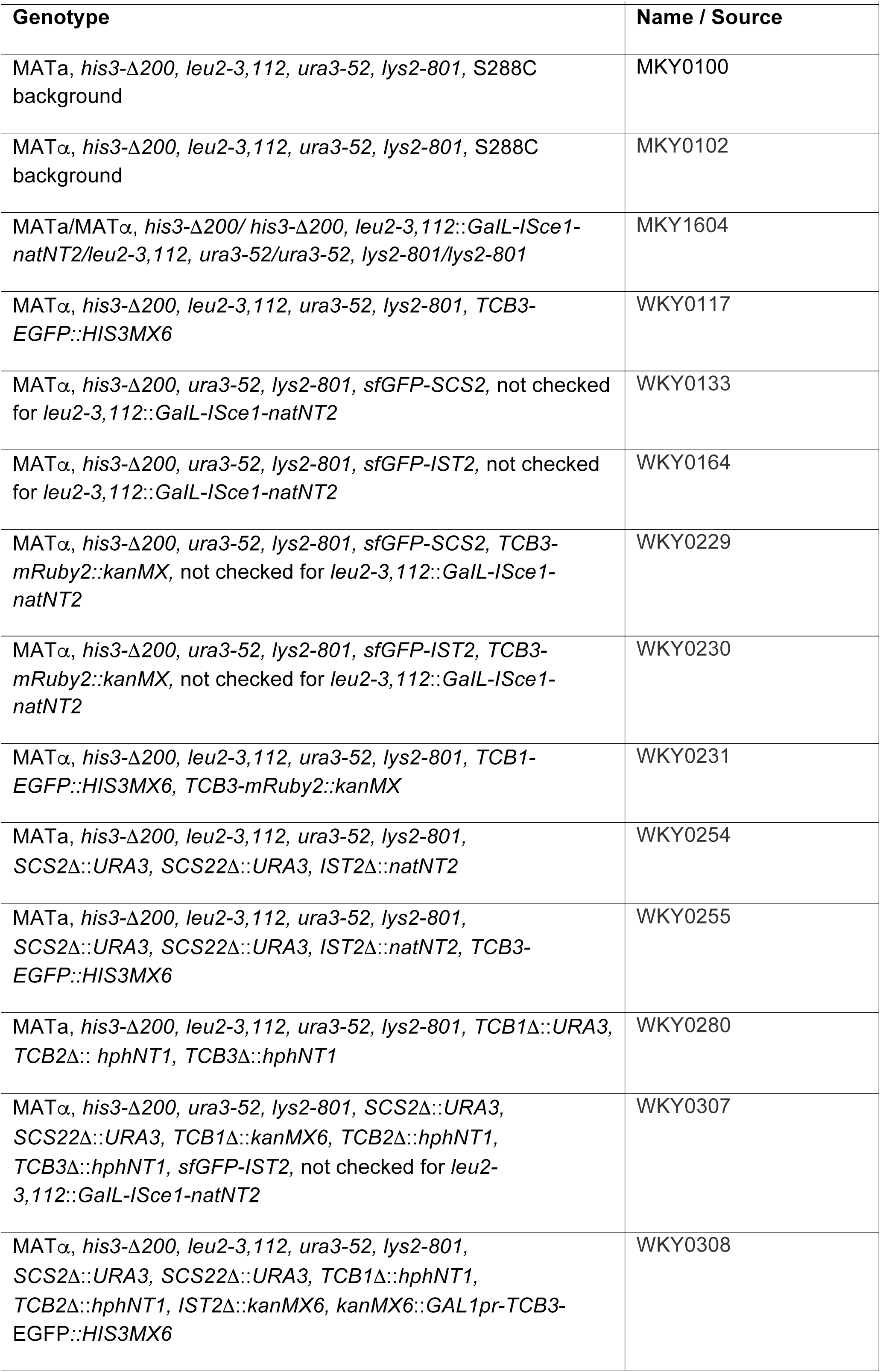
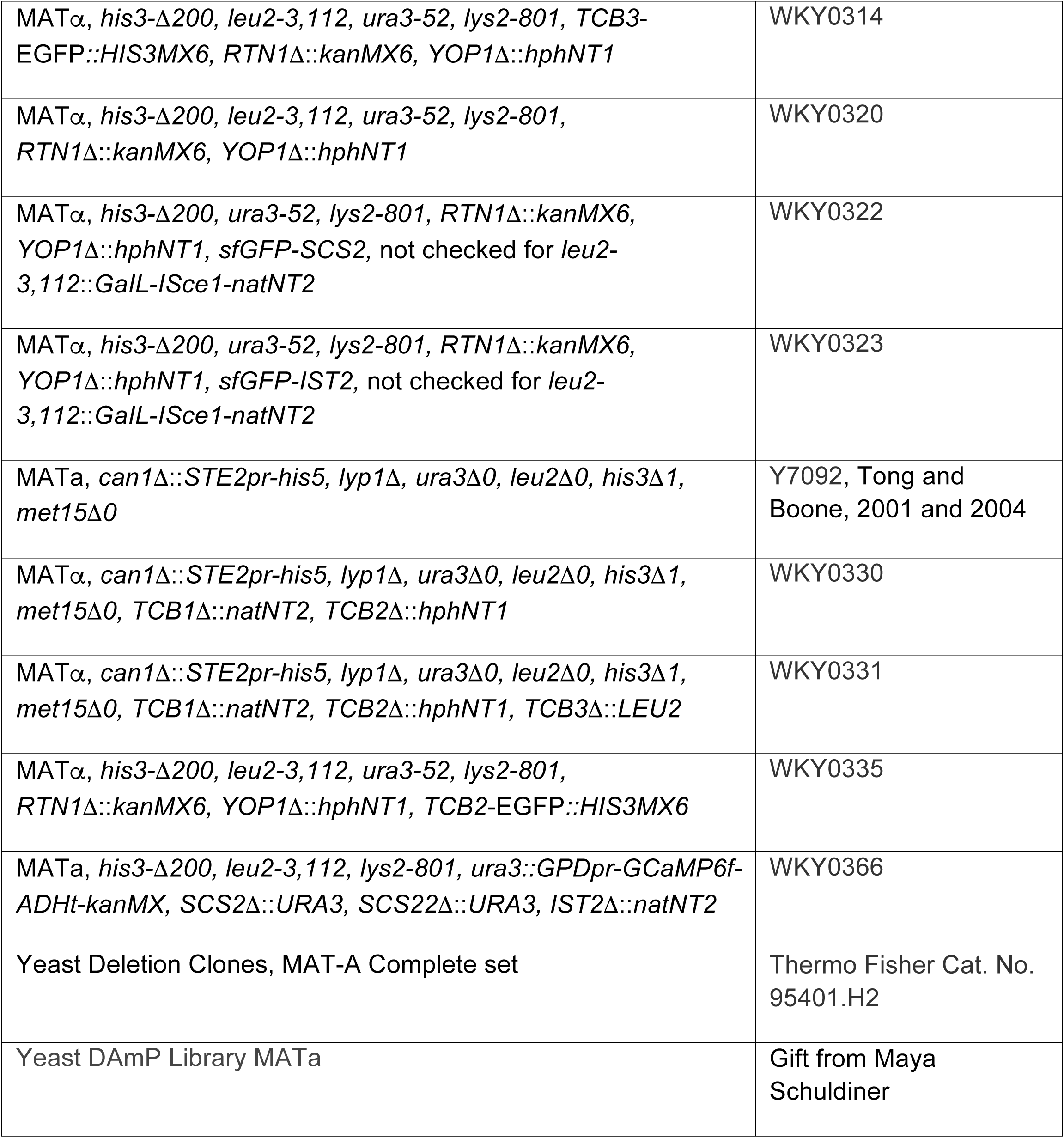
Yeast strains

